# Host-pathogen Coadaptation by running with PAR protein

**DOI:** 10.1101/2024.01.31.578330

**Authors:** Sucheng Zhu, Xinyi Zhou, Bin Qi

## Abstract

Host-pathogen coadaptation is mutually beneficial for the survival of both organisms. Embryo development is a crucial aspect of animal evolution as it ensures the continuation of their offspring. However, there is little understanding of the coadaptation mechanisms underlying embryo developmental factors in host-pathogen interactions. Here, by using a *C. elegans-P. aeruginosa* infection model, we demonstrate that PAR-5, essential for the establishment of anterior-posterior polarity within the single cell *C. elegans* zygote, acts as a mediator in regulating host-pathogen co-evolution. We discover that *Pseudomonas aeruginosa*-PA14 infection induces PAR-5 expression, which accelerates animal embryo development and ensures the continuity of their offspring. Moreover, PA14 stimulates host PAR-5 secretion, which reduces the virulence of PA14. Both functions of PAR-5 are beneficial for host survival. Meanwhile, pathogens utilize PAR-5 to inhibit host immunity/UPR^ER^ by bypassing the core PMK-1-mediated innate immunity and activating mTOR by interacting with LET-363, promoting pathogen spread. Therefore, our study uncovers an unexpected mechanism in host-pathogen co-evolution that targets a host embryo developmental factor, promoting coexistence and facilitating the adaptation of both hosts and pathogens.

## Introduction

Host-pathogen coevolution provides important insights into the intricate interplay between hosts and pathogens, which is crucial for understanding the emergence and spread of infectious diseases. Hosts develop multiple antibacterial mechanisms to detect and limit pathogen infections, while pathogens coevolve by employing various strategies to evade host cellular functions. This process accelerates relevant protein evolution (Nielsen, 2005). The host acquires resistance through the evolution of host genes under this arm race. The reproductive system is critical for animals since it ensures the continuity of their offspring. Many key factors regulating embryo development are conserved in animals (Goldstein and Macara, 2007). Moreover, evolutionary mutations of these factors are very fatal for animals. Thus, it is an ideal strategy for pathogens to evade the host immune response by hijacking these factors under continuous host-pathogen arm race.

Animals face a variety of challenges in their natural habitats, including limited food sources, high pathogen levels, and potential conflicts with other creatures. To survive and thrive in these environments, animals have developed both innate coping mechanisms and adaptive responses to stressors (McEwen and Wingfield, 2003). The reproductive system ensures the continuity of their offspring. One example of an adaptive response is the tendency for some animals to accelerate their offspring’s reproduction when facing severe environmental stress (Ding et al., 2023; Paxton et al., 2016), which can help them to maintain their existence. From the perspective of pathogen-host coevolution, pathogens may also hijack host factors involved in reproduction or embryo development for their infection. However, the importance of specific factors involved in the reproductive system for maintaining host-pathogen coevolution/co-adaptation and the underlying mechanism remain unclear. We hypothesize that pathogens may target host key embryo-developmental factors for their infection and host offspring production, ultimately benefiting host and pathogens for co-adaptation and co-survival during evolution.

## Results

### PAR-5 promotes fertilization under infection, which benefits to the host

To survive, animals must ensure the continuity of their life. Animals may adapt to the environment by accelerating reproduction and producing offspring in a stressful environment. To test this hypothesis, we used a *C. elegans-P. aeruginosa* infection model to analyze the number of fertilized eggs of wild-type animals (N2) exposed to either *E. coli-*OP50 (OP50) or *P. aeruginosa-*PA14 (PA14). We found that an increased number of zygotes during the process from the first fertilized egg formation in the body until it was expelled in animals exposed to PA14 compared to worms grown in OP50 (Figure 1A), suggesting that *C. elegans* accelerates egg formation to reproduce offspring during PA14 infection. A series of studies have revealed that the accelerated development of animals is closely related to the micronutrients, particularly Vitamin B12 (VB12), produced by specific bacterial strains (Bito et al., 2013; Watson et al., 2014) It has been found that the bacterial strain PA14 is capable of synthesizing Vitamin B12, unlike the *E. coli-*OP50 strain (Crespo et al., 2018; Watson et al., 2014). To investigate whether the acceleration of egg formation in worms during PA14 infection is due to the higher VB12 content in PA14, we conducted assays using *E. coli-*OP50 supplemented with VB12 as a control. The supplementation of OP50 with VB12 resulted in an increased number of fertilized eggs, similar to the observation in wild-type animals exposed to PA14 (Figure S1A). This suggests that worms may obtain a great amount of micronutrients, specifically Vitamin B12, from PA14, which accelerated eggs formation to reproduce offspring.

**Figure 1.**
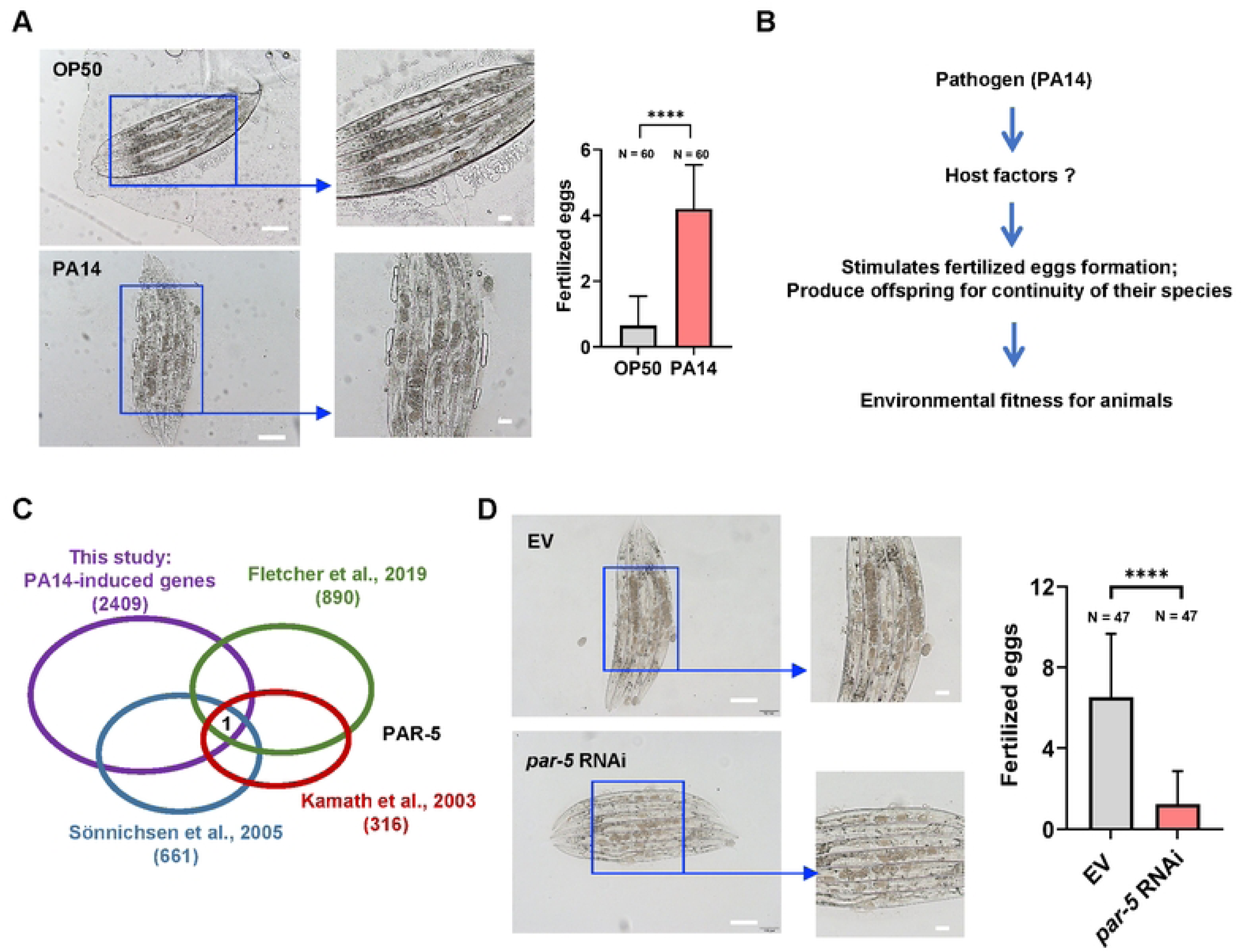
PAR-5 mediates fertilization, embryonic development and offspring production in *C. elegans* during pathogen infection. (A) Microscope images and bar graph showing PA14 stimulates fertilized eggs formation. L4 *C. elegans* were infected with PA14 for 7 h, then number of fertilized eggs were recorded during the process from the first fertilized egg formation in the body until it was expelled in animals. Scale bar, 100 μm. (B) Text illustration of the hypothesis that PA14 induces some host factors in *C. elegans* that contribute to accelerate egg-formation and offspring production for continuity of their species. (C) A Venn diagram comparing the overlap among embryogenesis-related genes from studies published by Kamath et al (Kamath et al., 2003) and Sönnichsen et al (Sönnichsen et al., 2005), genes up-regulated by PA14 infection (this study) and RNA-seq data sets from the study published by Fletcher et al (Fletcher et al., 2019) (see Table S2). (D) Microscope images and bar graph showing that formation of fertilized eggs inducing by PA14 was reduced in animals with *par-5* RNAi. L4 *C. elegans* were infected with PA14 for 18 h, then number of fertilized eggs were recorded. Scale bar, 100 μm. For all panels, N is the number of worms scored. Data are represented as mean ± SD. ∗∗∗∗p < 0.0001, ∗∗∗p < 0.001, ∗∗p < 0.01, ∗p < 0.05, ns, no significant difference. All data are representative of at least three independent experiments.

Previous research has shown that worms with defects in egg-laying often exhibit a phenomenon known as “worm bagging,” where the eggs of nematodes hatch inside the parent’s body during pathogen infections (Mosser et al., 2011). To rule out the differences of the fertilized eggs formation inside the insect’s body due to egg retention during pathogen infection, we analyzed the Bag phenotype of worms exposed to *E. coli*-OP50 or PA14, and found that nematodes could lay and ovulate eggs normally under PA14 conditions (Figure S1B), without forming any “worm bagging” (Figure S1C). This result suggests that the phenomenon of PA14 stimulating rapid egg production is not due to the occurrence of “worm bagging.”

Thus, we speculate that some host factors in *C. elegans* may be induced during PA14 infection and contribute to accelerate egg-formation and offspring production for adapting to the stressful environment (Figure 1B). To find host factors involving PA14-induced egg-formation, we first performed RNA-seq on wild-type (N2) animals exposed to OP50 or PA14 to identify genes that are upregulated upon PA14 infection. We found that 2409 genes were two-fold upregulated at 8 hours post-infection, compared to animals exposed to OP50 (Figure 1C). Specifically, only one shared gene (*par-5*) involved in embryogenesis, published by Kamath et al (Kamath et al., 2003) and Sönnichsen et al (Sönnichsen et al., 2005) , overlapped between our data sets and the RNA-seq data sets from the study published by Fletcher et al (Fletcher et al., 2019) (Figure 1C). PAR-5 is the *C. elegans* homolog of the mammalian 14-3-3 protein, which plays a crucial role in embryonic cell fates (Morton et al., 2002). Thus, we asked whether *par-5* also contributes to offspring formation in worms exposed to PA14. We analyzed the egg-laying rate and hatching rate of N2 animals with or without *par-5* RNAi and found that the egg-laying rate and hatching rate were decreased in animals treated with *par-5* RNAi (Figure S1D and Figure S1E), consistent with the previous conclusion that *par-5* mutation leads to embryonic lethality and sterility (Mikl and Cowan, 2014). Furthermore, PA14 induced fertilized eggs-formation is decreased in animals with *par-5* RNAi (Figure 1D). These results suggest that PAR-5 plays a key role in offspring production and the development of worms, and may explain why the expression of *par-5* is induced in *C. elegans* upon PA14 infection. Taken together, these data indicate that PAR-5 may be an important regulator that improves worm adaptability to *P. aeruginosa* infection by shortening the spawning period, thereby enabling offspring to reproduce early, which is critical for animal survival and reproduction in the face of pathogen infection.

### PA14 utilizes PAR-5 for infection, which is beneficial for the pathogens

By using *C. elegans-P. aeruginosa* interaction model, we have identified that host factor PAR-5 mediates fertilization and embryonic development of *C. elegans* to enhance their adaptability to the PA14 infection, which is beneficial for the animals (Figure 2A). From the perspective of host-pathogen co-evolution, we asked if the pathogen (PA14) utilizes the host factor PAR-5 to infect the host. We speculated that PA14 might suppress innate immunity by hijacking the host factor PAR-5 to infect nematodes, which is beneficial for the pathogens (Figure 2A).

**Figure 2.**
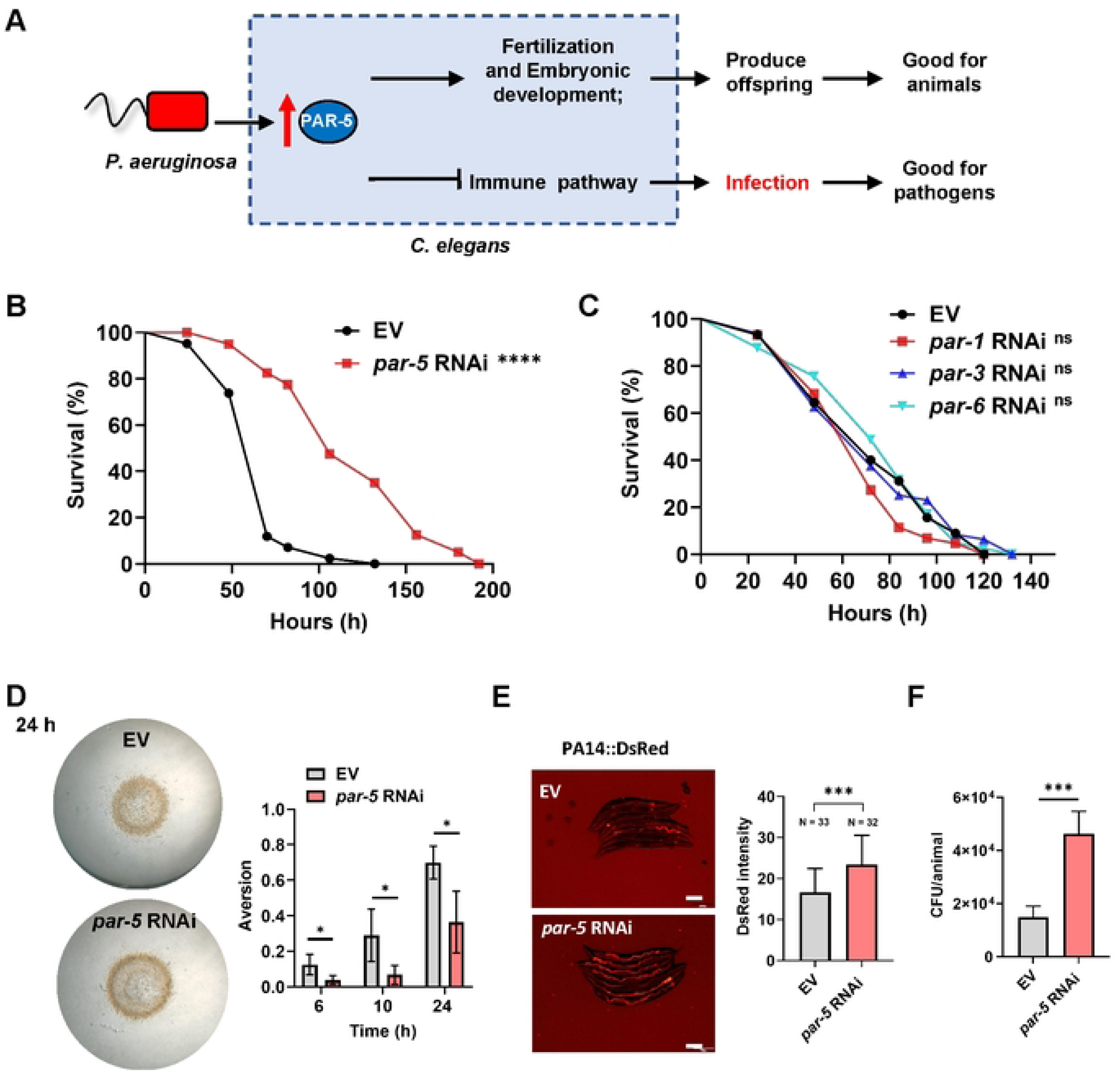
PA14 infects worms by hijacking host PAR-5. (A) Cartoon illustration of the hypothesis that PA14 induces host factor PAR-5 in *C*. *elegans* to enhance their environmental adaptation by increasing offspring production and development, while PA14 also hijacks host factor PAR-5 to suppress the innate immunity and infect nematodes. (B) Animals with *par-5* RNAi significantly enhances the resistance to PA14 infection. *n = 50* worms for each sample. (C) Knockdown of PAR protein families-related genes (*par-1, par-3* and *par-6*) by RNAi has no effect the survive of worms exposed to PA14. *n = 50* worms for each sample. (D) Microscope images and bar graph showing RNAi inhibition of *par-5* reduces PA14 lawn avoidance behavior in small lawn. More than 70 worms were used. (E) Representative images and quantification of PA14-dsRed colonization in animals grown on control RNAi bacteria or *par-5* RNAi bacteria to L4 stage followed by exposure to PA14-dsRed for 24 h at 20°C. Scale bar, 200 μm. (F) Numbers of colony-forming units of PA14 were measured in worms subjected to *par-5* RNAi. These results are mean ± SD of four independent experiments performed in triplicate For all panels, N is the number of worms scored. Data are represented as mean ± SD. ∗∗∗∗p < 0.0001, ∗∗∗p < 0.001, ∗∗p < 0.01, ∗p < 0.05, ns, no significant difference. All data are representative of at least three independent experiments. The exact *P* values of statistics for all survival assays are listed in Table S1.

To confirm this hypothesis, we performed survival assay in animals with or without *par-5* RNAi exposed to PA14 infection, and found that animals with *par-5* RNAi enhance the resistance to the PA14 infection (termed as <u>P</u>athogen <u>R</u>esistance <u>P</u>henotype: PRP) (Figure 2B). In *C. elegans*, the *ftt-2* gene is homologous to *par-5*, sharing approximately 86% homology in bases and 80% in amino acids (Wang and Shakes, 1997). PAR-5 controls asymmetries of other PARs proteins which plays a crucial role in the early events leading to polarization of the *C. elegans* zygote (Goldstein and Macara, 2007). To determine if the <u>P</u>athogen <u>R</u>esistance <u>P</u>henotype (PRP) was specific in animals with RNAi of *par-5*, we performed survival assays in animals with *ftt-2, par-1, par-3* and *par-6* RNAi. We found that animals with other *par* RNAi had no effect on the survival of worms exposed to PA14 infection (Figure 2C). However, animals with *ftt-2* RNAi were more sensitive to PA14 infection (Figure S2A). These results suggest that PA14 may infect animals by specifically targeting PAR-5, resulting in a Pathogen Resistance Phenotype (PRP) that is only observed in animals with *par-5* RNAi.

Several studies have shown that nematodes exhibit behavioral pathogen avoidance to improve their survival when infected with PA14 (Meisel and Kim, 2014; Reddy et al., 2009; Styer et al., 2008). To determine if the *par-5* RNAi affects PA14-avoidance behavior and causes the Pathogen Resistance Phenotype (PRP) phenotype, we exposed animals to PA14 lawns. We found that animals with *par-5* RNAi escaped slower from the bacterial lawn than vector control RNAi worms (Figure 2D), consistent with previous studies (Hong et al., 2021). This result suggests that the PRP phenotype caused by *par-5* RNAi is not related to PA14-avoidance behavior.

Pathogenic bacteria colonization in animal gut is considered a key factor in PA14-induced killing (Tan et al., 1999). A previous study showed that animals increase resistance to PA14 infection while reducing intestinal pathogenic bacterial colonization (Evans et al., 2008). To investigate if reduced intestinal colonization of PA14 (termed pathogen clearance) contributes to the Pathogen Resistance Phenotype (PRP) phenotype in animals with *par-5* RNAi, we analyzed bacterial colonization in animals exposed to PA14 expressing red fluorescent protein (PA14-dsRed) and OP50 expressing mCherry (OP50-mCherry), and found that colonization of PA14-dsRed (Figure 2E) and OP50-mCherry (Figure S2B) are increased in animals with *par-5* RNAi. .Furthermore, *par-5* RNAi markedly increased the colony forming units (CFU) of PA14 (Figure 2F) or OP50 (Figure S2C) in worms. Thus, these results suggest that PAR-5 negatively impacts bacterial colonization. To rule out changes in colonization due to differences in bacterial consumption, we measured the pumping rate of worms exposed to *E. coli*-OP50 or PA14. The experiment showed no difference in pumping rate between wild-type and *par-5* RNAi worms (Figure S2D), indicating that PAR-5 did not affect nematode feeding on bacteria. We also monitored lumen morphogenesis using ERM-1::GFP reporter animals (Göbel et al., 2004) and found that *par-5* RNAi caused bloating of the intestinal lumen (Figure S2E), suggesting that bacterial accumulation may result in lumen bloating.

Taken together, these results suggest that PAR-5 is critical target for PA14 infection, and the Pathogen Resistance Phenotype (PRP) caused by *par-5* RNAi is not related to the PA14-avoidance behavior (Figure 2D) or pathogen clearance (Figure 2E and 2F), implicating that PA14 may hijack the PAR-5 to inhibit host immune system for infection.

### Intestinal PAR-5 is targeted by PA14 for infection

To examine the expression pattern of PAR-5, we used a transgenic worm (*par-5p::gfp::par-5*) for observation. We found strong PAR-5 expression in the intestine, head neurons and pharynx (Figure 3A). Additionally, we discovered that *par-5* RNAi in intestine enhanced the resistance of worms to PA14 infection (Figure 3B), but not in neurons, muscle or hypodermis (Figure 3C-E), suggesting that PAR-5 is functional in intestine for the Pathogen Resistance Phenotype (PRP) in animals with *par-5* RNAi. Consistent with this idea, the transgenic worms with intestine-specific overexpression of *par-5* were sensitive to PA14 compared to the control animals (Figure S3A). Meanwhile, we silenced *par-5* in the intestinal *par-5* overexpression animals and found that these worms had greater resistance to the PA14 infection compared to the control groups (Figure S3B). These results indicate that PAR-5 functions in the intestine in response to PA14 infection.

**Figure 3.**
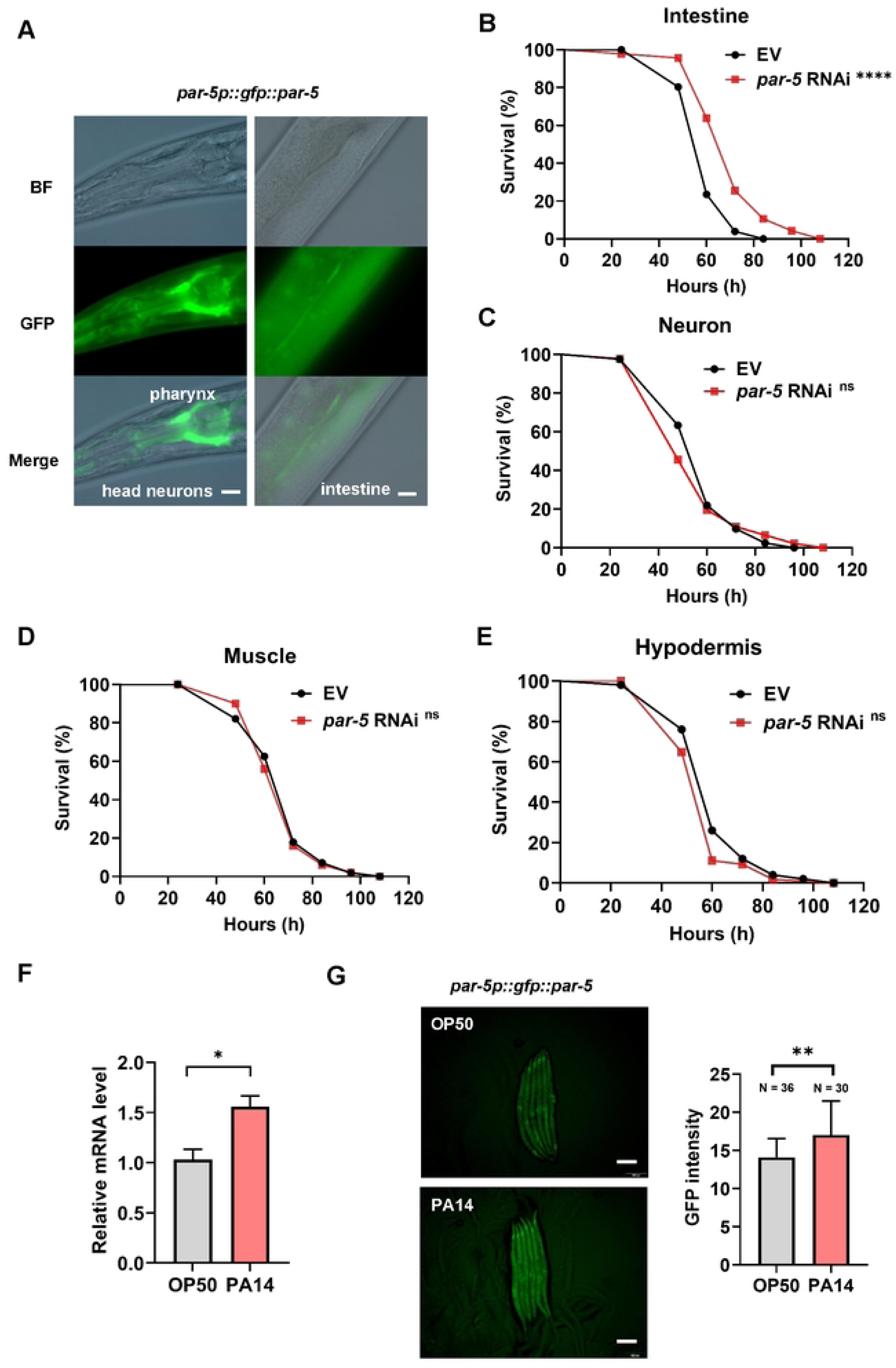
PA14 induces PAR-5 expression and intestinal PAR-5 is targeted by PA14. (A) Bright-field and fluorescence images showing the expression of a single-copy insertion of the *par-5p::gfp::par-5* reporter in the intestine, head neurons and pharynx. Scale bar, 10 μm. (B) Animals with intestinal *par-5* RNAi significantly enhances the resistance to PA14 infection. *n = 55* worms for each sample (C-E) Survival assay of animals (tissue specific RNAi strains) with *par-5* RNAi in neuron (C) (*n = 55*), muscle (D) (*n = 50*) and hypodermis (E) (*n = 60*) under PA14 infection. (F) Results of qRT-PCR analysis showing the expression of *par-5* in wild-type (N2) animals exposed to *E. coli* OP50 or PA14. (G) Expression of *gfp::par-5* is up-regulated in WT worms exposed to PA14 for 48h at 20°C. Scale bar, 100 μm. For all panels, N is the number of worms scored. Data are represented as mean ± SD. ∗∗∗∗p < 0.0001, ∗∗∗p < 0.001, ∗∗p < 0.01, ∗p < 0.05, ns, no significant difference. All data are representative of at least three independent experiments. The exact *P* values of statistics for all survival assays are listed in Table S1.

To confirm that the expression of *par-5* is induced upon PA14 infection, we used quantitative real-time PCR (qRT-PCR) to measure *par-5* transcript levels in worms exposed to PA14. The results showed that the expression of *par-5* was significantly upregulated in animals exposed to PA14 for 12 hours, compared to worms grown in *E. coli* OP50 (Figure 3F). Moreover, transgenic animals expressing *par-5p::gfp::par-5* showed increased GFP expression after PA14 infection (Figure 3G), indicating that PAR-5 is induced by PA14.

Furthermore, we investigated whether PA14 activates the expression of *par-5* through its virulence factors. We screened PA14 mutants (Figure S3C) that lacked key virulence factors involved in host infection (Qin et al., 2022). Our results showed that worms exposed to PA14 mutants *pqsF*, *lasR*, *mvfR* and *gacA* had significantly decreased *par-5::gfp* expression compared to those exposed to wild type PA14 (Figure S3D). These findings suggest that the induction of *par-5* expression by PA14 may depend on its virulence factors.

### PA14 inhibits host immune and UPR^ER^ through PAR-5 for infection

To understand the mechanism underlying PAR-5’s involvement in pathogen response, we used RNA-seq to examine gene expression in N2 animals with or without *par-5* RNAi during PA14 infection. Of the 12,399 genes quantified, 4344 were up-regulated and 3697 were down-regulated by at least 2-fold in animals with *par-5* RNAi (Figure S4A and Table S3). GO enrichment analysis of the 4344 up-regulated genes identified 16 significantly enriched biological processes, including innate immune response, defense response to Gram-negative bacterium, cellular response to DNA damage stimulus, and endoplasmic reticulum unfolded protein response (UPR^ER^) (Figure 4A and Table S4). These findings suggest that the activation of these pathways may contribute to the Pathogen Resistance Phenotype (PRP) in animals with *par-5* RNAi.

**Figure 4.**
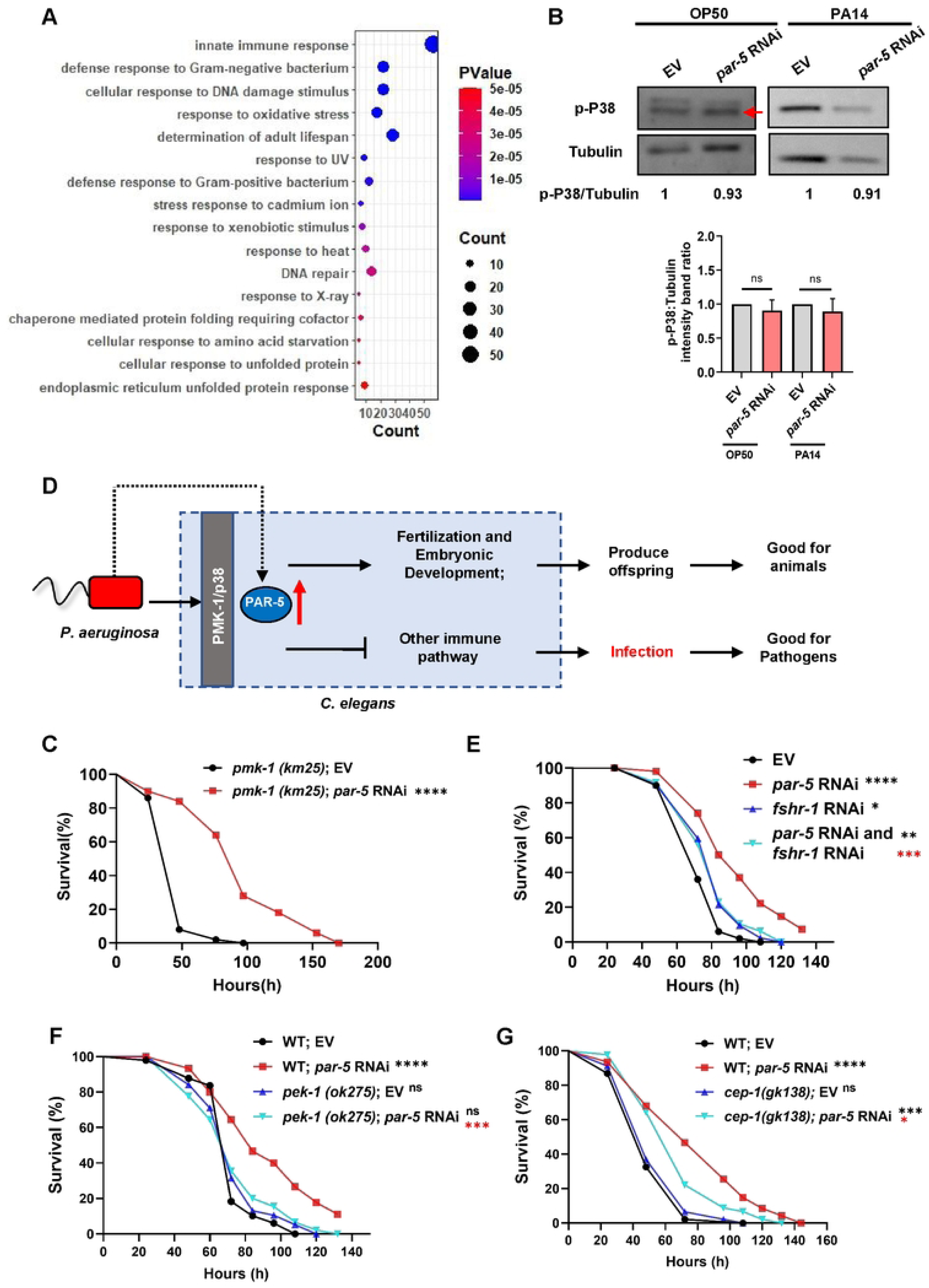
Knock-down PAR-5 activates host immune and stress response for anti-infection. (A) GO enrichment of up-regulated gene in animals with *par-5* RNAi under PA14 infection. The up-regulated genes are shown in Figure S4A and Table S4. (B) Western blot showing the level of p-PMK-1 in animals grown on *E. coli* OP50 or PA14 fed with control RNAi bacteria or *par-5* RNAi bacteria, and calculation of their relative densitometric ratios to p-P38/Tubulin shown in below. Data are shown as mean ± SD of more than three independent experiments. (C) *par-5* RNAi significantly enhances resistance to PA14 infection in *pmk-1 (km25)* mutant. Three independent experiments were performed. *n = 55* worms for each sample. (D) Cartoon illustration of the hypothesis that PA-induced PAR-5 increases offspring production and development in *C. elegans* for environmental adaptation (good for animal), while PA14 also hijacks PAR-5 to suppress the innate immunity and infect nematodes (good for pathogen). (E) *fshr-1* RNAi significantly reduces survival of worms with *par-5* RNAi upon PA14 infection. *n = 55* worms for each sample. (F) *pek-1* mutation significantly reduces survival of worms with *par-5* RNAi upon PA14 infection. *n = 50* worms for each sample. (G) *cep-1* mutation significantly reduces survival of worms with *par-5* RNAi upon PA14 infection. *n = 50* worms for each sample Data are represented as mean ± SD. ∗∗∗∗p < 0.0001, ∗∗∗p < 0.001, ∗∗p < 0.01, ∗p < 0.05, ns, no significant difference. All data are representative of at least three independent experiments. The exact *P* values of statistics for all survival assays are listed in Table S1.

The p38 mitogen-activated protein kinase (MAPK) pathway plays a crucial role in innate immune defenses in animals (Arthur and Ley, 2013). In *C. elegans*, the PMK-1/p38 pathway regulates the expression of immune effectors and is essential for survival during pathogen infection (Kim et al., 2002; Troemel et al., 2006). To determine the involvement of PAR-5 in the activation of the p38/PMK-1 innate immune pathway, we conducted experiments to test whether *par-5* RNAi triggers this pathway. Firstly, we found that *par-5* RNAi did not alter the expression of GFP in transgenic animals expressing *sysm-1p::gfp* or *irg-5p::gfp*, which are downstream effector molecules of the PMK-1 signaling pathway (Bolz et al., 2010; Soltanmohammadi et al., 2022) (Figure S4B). Secondly, immunoblotting against the activated form of PMK-1 showed that the levels of activated PMK-1 (p-PMK-1) in worms with *par-5* RNAi were similar to those observed in wild-type animals exposed to *E. coli* OP50 or PA14 (Figure 4B). Furthermore, *par-5* RNAi resulted in an increased resistance to PA14 infection in *pmk-1 (km25)* mutant compared to the vector control RNAi (Figure 4C). Based on these results, we conclude that PAR-5 is not involved in activating the PMK-1/p38 innate immune pathway in response to PA14 infection. Instead, knocking down of PAR-5 likely mediates worm defense against this pathogen through a PMK-1/p38-independent pathway.

PMK-1-mediated innate immunity is a major defense in nematodes. How does RNAi of *par-5* allow animals to resist pathogen infection through a PMK-1-independent pathway? Given that RNA-seq data showed numerous upregulated genes in animals with *par-5* RNAi involved in innate immunity and stress response (Figure 4A), we speculate that pathogens may bypass the core PMK-1-mediated innate immune by hijacking PAR-5 to suppress other immune pathways or stress response pathway to achieve infection, which is beneficial for the pathogens (Figure 4D).

Our RNA-seq showed that key immune and stress response factors are induced in animals with *par-5* RNAi (Figure S4C), including i) the G protein-coupled receptor FSHR-1 which is required for the *C. elegans* innate immune response (Powell et al., 2009); ii) the IRE-1-XBP-1 and PEK-1 which are required for maintaining ER homeostasis in animals under physiological ER stress (Richardson et al., 2011); iii) the p53/CEP-1 which required for activation of immunity against *S. enterica (Fuhrman et al., 2009).* Therefore, we asked if Pathogen Resistance Phenotype (PRP) in animals with *par-5* RNAi depends on the these immune and stress response factors inducing by *par-5* knock-down. Our results show that Pathogen Resistance Phenotype (PRP) in animals with *par-5* RNAi was all most suppressed by knocking-down of *fshr-1* (Figure 4E)*, or mutation of pek-1* (Figure 4F) and *cep-1* (Figure 4G), but not *xbp-1* (Figure S4D). Moreover, we found that UPR^ER^ was activated in animals with *par-5* RNAi by analyzing the expression level of *hsp-4p::gfp* (Figure S4E), which is independent of XBP-1 (Figure S4F).

Taken together, these results suggested that pathogens bypass the core PMK-1-mediated innate immune by hijacking PAR-5 to suppress FSHR-1/PEK-1/CEP-1-mediated immune or stress response to achieve infection, which is beneficial for the pathogens.

### Pathogens activate mTOR by targeting PAR-5-LET-363 complex for its infection

In order to further investigate the mechanism by which PAR-5 mediates infection by PA14, we performed immunoprecipitation (IP) with PAR-5::GFP protein against total isolated protein from worms exposed to PA14 (Figure 5A) and conducted mass spectrometry (MS) to identify proteins that potentially interact with PAR-5. The MS analysis revealed that components related to TOR signaling, including *F39C12.1*, *let-363*, and *rict-1*, interact with PAR-5 (Figure 5B). If the pathogens also utilize PAR-5-interacitng proteins for its infection, animals with knock-downing interaction proteins may also show Pathogen Resistance Phenotype (PRP), which is similar with *par-5* RNAi.

**Figure 5.**
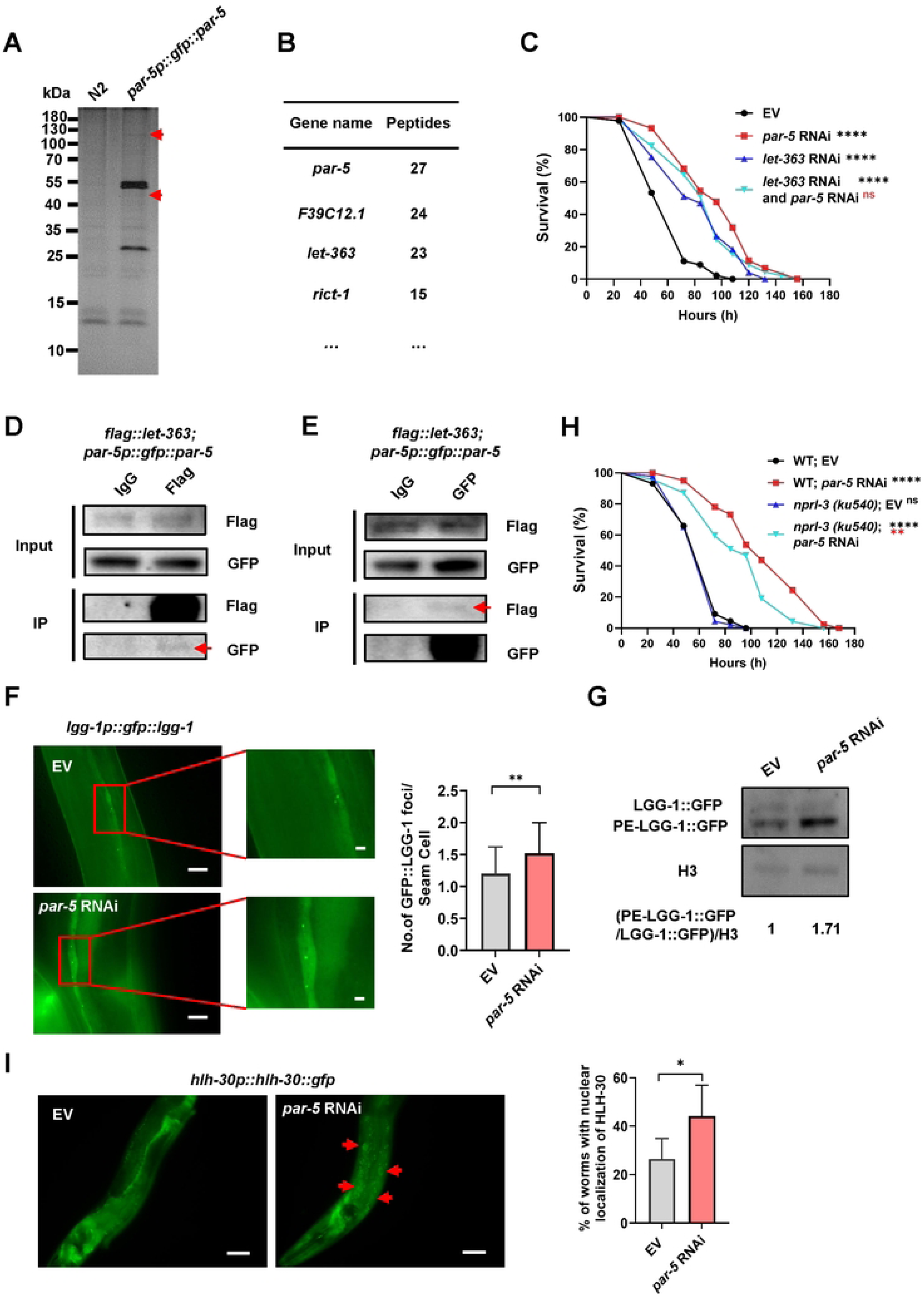
PAR-5 interacts with LET-363, which was targeted by PA14 for its infection. (A and B) Immunoprecipitation of PAR-5::GFP from control (N2) and *par-5p::gfp::par-5* transgenic animals with PA14 infection was shown by sliver-stained gel (A), and list of the PAR-5 interaction proteins identified by mass spectrometry (B). To identify the proteins interacting with PAR-5 under infection conditions, transgenic animals (*par-5p::gfp::par-5*) and wild-type N2 animals (as a control to remove unspecific binding proteins) were infected with PA14. After infection, total proteins from both transgenic and control animals were isolated. Then, GFP-antibody was added into both groups to pull down proteins interacting with PAR-5. The wild-type N2 animals as a control to remove proteins that unspecific bind to the antibody. (C) Animals with *let-363* RNAi enhanced the resistance to PA14 infection. Pathogen Resistance Phenotype (PRP) in animals with *par-5* RNAi was not enhanced by knock-downing *let-363. n = 50* worms for each sample (D and E) An *in vivo* IP experiment validates the interaction between PAR-5 and LET-363 using transgenic animals expressing *flag::let-363* + *par-5p::gfp::par-5*. For Co-IP, lysate of transgenic worms was incubated with anti-FLAG (D) or anti-GFP (E) beads (the IgG beads as negative control) for pulling down FLAG-LET-363-binding or GFP-PAR-5-binding proteins. FLAG-LET-363-binding or GFP-PAR-5-binding proteins were eluted and detected by western blot (anti-GFP or anti-FLAG). (F) Representative images of autophagosomes (GFP::LGG-1 puncta) in the seam cells of worms with *par-5* RNAi. The average number of GFP::LGG-1 puncta in each seam cell were counted (Right). Scale bar, 10 μm. *n ≥ 10* worms per condition. (G) Western blot showing the level of PE-conjugated LGG-1–GFP and LGG-1–GFP in animals fed with control RNAi bacteria or *par-5* RNAi bacteria. The blot is typical of three independent experiments. (H) *nprl-3* mutation reduces survival of worms with *par-5* RNAi upon PA14 infection. *n = 50* worms for each sample. (I) *par-5* RNAi promotes the nuclear localization of HLH-30::GFP. (n = 3 independent biological replicates, about 20 animals were examined for each replicate). Arrows indicate nuclear HLH-30::GFP signals. Scale bars: 20 μm. Data are represented as mean ± SD. ∗∗∗∗p < 0.0001, ∗∗∗p < 0.001, ∗∗p < 0.01, ∗p < 0.05, ns, no significant difference. All data are representative of at least three independent experiments. The exact *P* values of statistics for all survival assays are listed in Table S1.

Firstly, we found that animals with RNAi of *let-363* (Figure 5C), but not *F39C12.1* and *rict-1* (Figure S5A), significantly enhanced the resistance to PA14 infection. Secondly, Pathogen Resistance Phenotype (PRP) in animals with *par-5* RNAi was not enhanced by knock-downing *let-363* (Figure 5C), suggesting that PAR-5 and LET-363 are in the same pathway which was targeted by PA14 for infection. Thirdly, we generated transgenic animals expressing *flag::let-363* in *par-5p::gfp::par-5* animals to confirm the interaction between PAR-5 and LET-363 *in vivo*. We found that FLAG::LET-363 pull-down identified GFP::PAR-5 as its interacting partner (Figure 5D), and vice versa (Figure 5E), indicating that PAR-5 interacts with LET-363 *in vivo*. Together, our results suggest that PA14 infect animals through PAR-5-LET-363 complex.

mTOR (LET-363) activates protein synthesis and inhibits autophagy for regulating cellular progress(Blackwell et al., 2019). Our above data show PA14 utilizes host PAR-5-LET-363 (mTOR) for infection. It is possible that pathogens infect worms through activation of mTOR by utilizing PAR-5. If this hypothesis is right, knocking down *par-5* could inhibit mTOR in animals with infection. Firstly, the protein level of LET-363 was decreased in *par-5* RNAi animals, indicating that PAR-5 positively regulates the expression of LET-363 (Figure S5B). Secondly, RNA-seq data showed that numerous gens involved in translation were down-regulated in animals with *par-5* RNAi under infection (Figure S5C and Table S5), which include ribosomal protein-related *rpl* genes and *rps* genes (Figure S5D and Figure S5E), suggesting that PAR-5 activates proteins synthesis in infection. Thirdly, we measured TORC1 activity by utilizing an established autophagy marker (LGG-1::GFP puncta) known to be repressed by TORC1 (Robida-Stubbs et al., 2012), and found that LGG-1::GFP puncta in seam cells is increased in animals with *par-5* RNAi (Figure 5F). Additionally, we also detected the ratio of phosphatidylethanolamine (PE)-conjugated LGG-1 (PE-LGG-1::GFP) to LGG-1 using Western blotting (Kang et al., 2007) to confirm the autophagy, and found that a significant increase in PE-LGG-1-GFP in *par-5* RNAi worms (Figure 5G). Fourth, Pathogen Resistance Phenotype (PRP) in animals with *par-5* RNAi was abolished by knock-out *nprl-3* which is a negative regulator of TORC-1 in yeast, worms and mammalian (Bar-Peled et al., 2013; Zhu et al., 2013) (Figure 5H). Finally, *par-5* RNAi resulted in significant nuclear translocation of HLH-30::GFP (Figure I) that can translocate to the nucleus to activate autophagy-related genes in response to inhibition of TOR signaling (Lapierre et al., 2013), indicating that *par-5* RNAi results in activation of HLH-30 during PA infection thereby leading to the activation of autophagy. Taken together, these data suggest that animals with *par-5* RNAi increased pathogen resistance by inhibiting mTOR which in turn inhibition translation and activation autophagy under infection condition. Our results indicated that pathogens activate mTOR by targeting PAR-5-LET-363 complex for its successful infection, which is benefit for pathogens.

### Secreted PAR-5 decrease the virulence of PA14 by interacting with virulence factors

During an infection by pathogens, *C. elegans* activates its immune signaling pathways to produce antibacterial proteins such as antimicrobial peptides, lectins, and lysozyme, which interact with and kill pathogenic bacteria (Mallo et al., 2002). In response to PA14 infection, host PAR-5 proteins are also induced (Figure 3G). We hypothesized that PA14-induced-PAR-5 could be secreted from worms and affect PA14 toxicity.

We cultured L1 animals with and without PA14 in liquid and collected the supernatant. We detected GFP::PAR-5 in the supernatant treated with PA14, but not under normal conditions, from *par-5p::gfp::par-5* animals (Figure 6A). This suggests that PA14 can induce PAR-5 secretion in worms. To determine the interaction between PAR-5 and PA14, we used PA14 to pull down His-tagged PAR-5 protein purified from *E. coli* BL21 transfected with *his::par-5* (Figure S6A). We found that His-PAR-5 directly binds to PA14 (Figure 6B). Interestingly, we also observed that peptidoglycan (PGN), an important structural element in the cell wall of bacteria, directly binds to His-PAR-5 (Figure S6B). This suggests that PAR-5 may interact with PA14 by binding to PGN in the cell wall.

**Figure 6.**
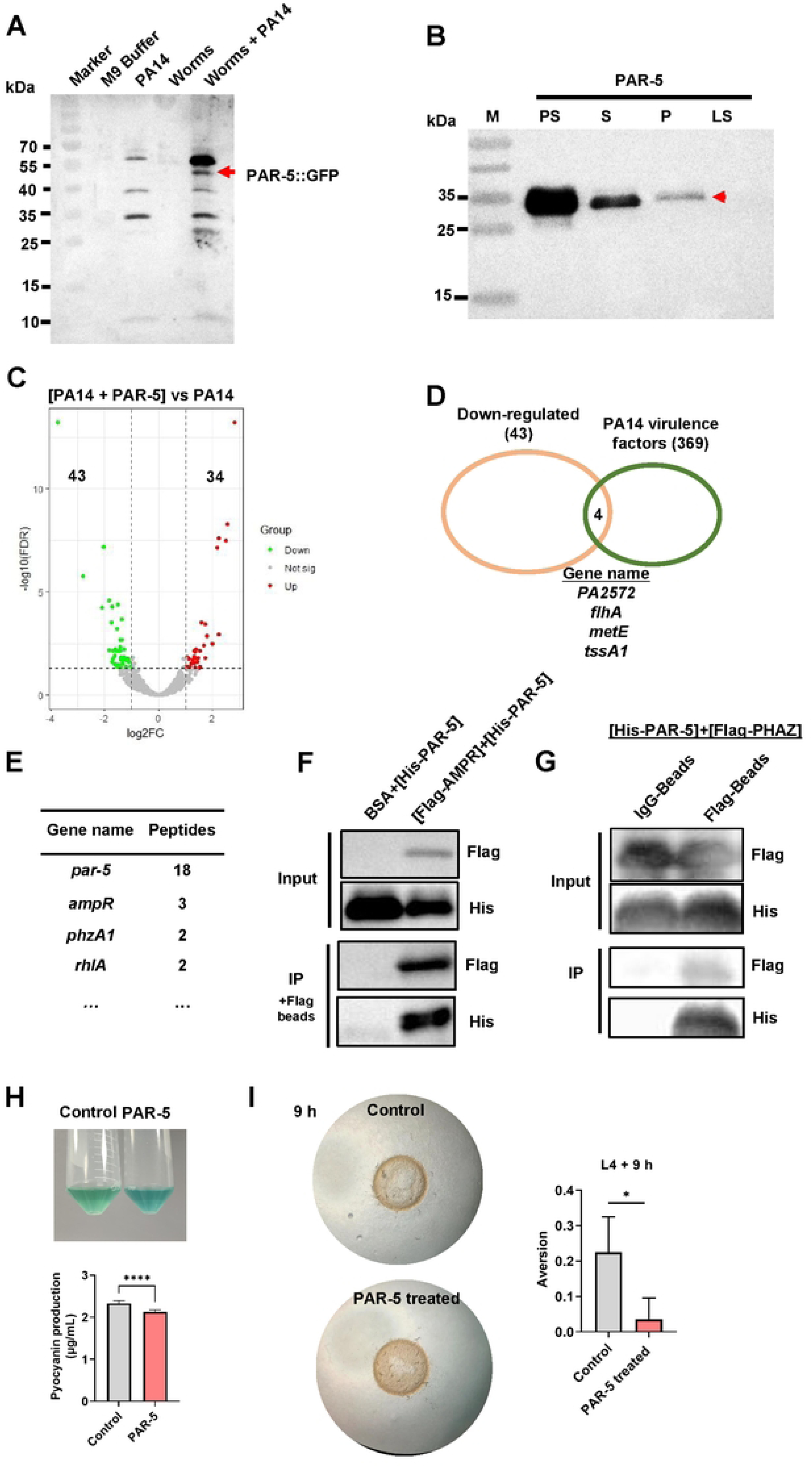
PAR-5 reduces virulence of PA14 by interacting with virulence factors. (A) Western blot showing that the secreted PAR-5::GFP was identified in the supernatant of liquid culture system which culture L1 *par-5p::gfp::par-5* transgenic animals with or without PA14. (B) *In vitro* bacterial binding assay shows that PA14 binds PAR-5 protein purified from *E. coli* BL21 transfected with *his::par-5*. PA14 was incubated with His-PAR-5 protein, the PA14 pellet was washed 3 times after incubation. Proteins associated with the washed pellet (P) and the incubated supernatant (S, LS) were detected by Western blot. M: marker; PS: purified PAR-5 protein; P: pellet; S: incubation supernatant; LS: supernatant of the last wash. (C) Volcano plot of transcripts corresponding to differentially expressed genes in PA14 treated with or without PAR-5 protein (see Table S6). Green and red colored dots indicate the differentially expressed genes that are increased or decreased, respectively. (D) A Venn diagram comparing the overlap of down-regulated genes and PA14 virulence-related genes (see Table S7). (E) List of the PAR-5 interaction proteins from the PA14 identified by mass spectrometry. (F) An *in vitro* IP experiment validates the interaction between PAR-5 and AMPR using Flag-AMPR protein from the lysate of *E. coli* BL21 (BSA protein as negative control) to pull down His-PAR-5 protein. (G) An *in vitro* IP experiment validates the interaction between PAR-5 and PHAZ using purified His-PAR-5 protein and *E. coli* BL21 expressing recombinant Flag-PHAZ protein. For Co-IP, the mixture of lysate of *E. coli* BL21 and His-PAR-5 protein was incubated with anti-FLAG beads (the IgG beads as negative control) for pulling down Flag-PHAZ-binding proteins. Flag-PHAZ-binding proteins were eluted and detected by western blot (anti-His or Anti-FLAG). (H) The images showing PAR-5 was added to ∼10^4^ CFU of log-phase PA14 for 8 hours, which changes the physiological behavior of PA14. The bar graph show pyocyanin production measured at OD = 695 nm in cell lysate prepared from PAR-5-treated PA14. (I) Microscope images and bar graph showing pathogen-induced avoidance was inhibited by PAR-5-treated PA14 lawn. More than 50 worms were used Data are represented as mean ± SD. ∗∗∗∗p < 0.0001, ∗∗∗p < 0.001, ∗∗p < 0.01, ∗p < 0.05, ns. All data are representative of at least three independent experiments.

**Figure 7.**
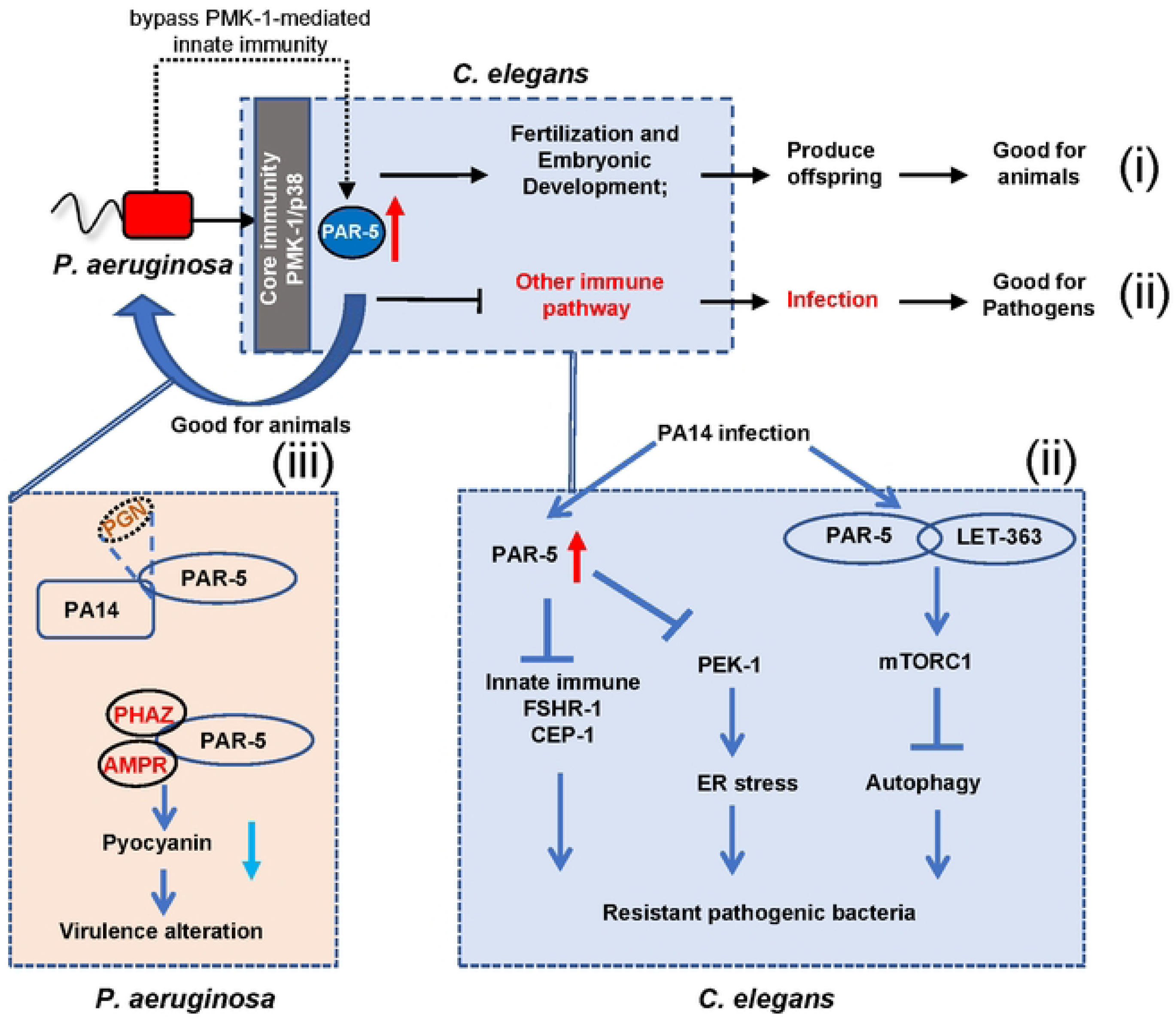
A model showing a critical role of host factor PAR-5 in regulating the co-adaptation between *C. elegans* and *P. aeruginosa* (i) Host factor PAR-5 is induced by virulence factors during PA14 infection, and contributes to offspring production of worms for environmental adaptation, which is beneficial for the host. (ii) PA14 inhibits host immunity/UPR^ER^ and activate mTOR through hijacking PAR-5, which is beneficial for the pathogens. (iii) PAR-5 is secreted from the animals under PA14 infection. Secreted PAR-5 reduces the virulence of PA14 by interacting with virulence factors, which is beneficial for the host. Host and pathogen co-existence and co-evolution by competing for PAR-5.

The direct interaction between PAR-5 and PA14 led us to hypothesize that PAR-5 may affect the physiological behavior of PA14. We tested the bactericidal activity of PAR-5 and found that it did not influence the viability of either enteric commensal (*E. coli* OP50) or pathogenic bacteria (PA14), even at high micromolar concentrations of PAR-5 (Figure S6C), although it can slightly affect the growth of *E. coli* OP50 at high concentrations (Figure S6C). These results indicates that PAR-5 is not a bactericidal protein.

To investigate whether PAR-5 affects the physiological state of PA14 by interacting with it, we used RNA-seq to profile gene expression in PA14 treated with or without PAR-5 protein. We found that 34 genes were upregulated and 43 genes were downregulated at least 2-fold in PAR-5-treated PA14 relative to untreated group (Figure 6C and Table S6). By overlapping the down-regulated genes with PA-related virulence factors, we found 4 genes (about 10% of down-regulated genes) that have been reported to be related to the virulence of PA (Figure 6D) (Qin et al., 2022). This suggests that PAR-5 may affect the virulence of PA14.

We also performed immunoprecipitation (IP) with purified His-PAR-5 against total isolated protein from PA14 (Figure S6D) and conducted mass spectrometry (MS) to identify proteins that potentially interact with PAR-5. The MS analysis revealed that 3 virulence factors, including *ampR*, *phzA1*, and *rhlA* (Azam and Khan, 2019), interact with PAR-5 (Figure 6E). To confirm this Co-IP result, we used purified His-PAR-5 protein to pull down Flag-tagged virulence-related proteins purified from *E. coli* BL21 expressing recombinant virulence proteins. We found that His-PAR-5 was identified by FLAG-AMPR pull-down and by FLAG-PHZA1 pull-down as their interacting partners (Figure 6F and 6G). This suggests that PAR-5 may affect the virulence of PA14 by interacting with AMPR and PHZA1.

To confirm if PAR-5 affects the virulence of PA14, we analyzed production of pyocyanin (an important virulence factor for PA14) (Lau et al., 2004) in PA14 treated with purified PAR-5 protein and found that PAR-5 inhibited pyocyanin production *in vivo* (Figure 6H). In addition, we also observed that pathogen-induced avoidance was inhibited in PAR-5-treated PA14 lawn (Figure 6I), indicating that PAR-5 may inhibit the virulence of PA14. Taken together, these results indicated that the host factor PAR-5 is induced by PA-14 to secrete out of the cells and may interact with virulence factors, including AMPR and PHZA1, to affect the virulence of PA14.

## Discussion

Here we found that co-evolutionary arm race between pathogen and host was mediated by the host embryo-developmental-factor, PAR-5. PA14 infection induces PAR-5 expression and secretion that promotes offspring production in worms and reduces the virulence of PA14, which is beneficial for the host survive. However, pathogens hijack host PAR-5 to inhibit host immunity/UPR^ER^ and activate mTOR for its spread, which is beneficial for the pathogens. Our findings show the critical role of host PAR-5 in regulating host-pathogen coadaptation.

The partitioning defective (PAR) genes were first identified in *C. elegans* but subsequently found in other organisms (Kemphues et al., 1988; Ohno, 2001), and play a crucial role in the establishment and maintenance of cell polarity by the zygote (Suzuki and Ohno, 2006). In *C. elegans*, PAR-related genes encode six different proteins (PAR-1, PAR-2, PAR-3, PAR-4, PAR-5 and PAR-6) and affect the asymmetric cell division in the worm zygote (Lang and Munro, 2017). Loss of any of the six PAR genes leads to loss of polarity, subsequent abnormal symmetric cell division, and ultimately embryonic lethality (Kemphues and Strome, 1997). However, besides PAR proteins function in maintaining embryo development, it is still not clear whether PAR proteins act in the regulating of host-pathogen coevolution. We found that PA14 only hijacks PAR-5, but not other PAR proteins, to inhibit host immunity/UPR^ER^ and activate mTOR for its infection. In animals, PAR-5 induced by the pathogens promotes offspring production in worms, which is critical for animal survival by continuity of their offspring.

Why do pathogens only targets PAR-5, but not other PARs? PAR-5 is a member of the 14-3-3 family of proteins, which controls asymmetries of the other PARs (Goldstein and Macara, 2007; Macara, 2004; Morton et al., 2002). It is possible that targeting PAR-5 by pathogens may promote embryo development and inhibit immune system in animals more efficiently, which benefits both pathogens and host. Unexpectedly, we also found that PAR-5 like antimicrobial peptides can be secreted out from worms after infection and decrease the virulence of PA14 by binding with virulence factors. In plant, 14-3-3 proteins have been found to interact with components of immune system to regulating immunity against pathogens (Konagaya et al., 2004). On the other hand, pathogens secrete effector proteins to target the 14-3-3 proteins to infect host (Giska et al., 2013; Kim et al., 2009; Taylor et al., 2012) , implying that 14-3-3 proteins might act as an important component in regulating the pathogen-host interaction. Therefore, we discovered an unexpected but logical role of PAR-5 in balancing the host-pathogen existence, which is important for their coadaptation.

Finally, our study points out that hijacking host embryo developmental factors for infection is an ideal strategy for pathogens, as evolutionary mutations of these factors are very fatal for animals. At this arm race, host also develop the strategy to use these factors to promotes offspring production and anti-infection for their survive. We propose that embryo developmental factors may act as mediator in maintaining host-pathogen coevolution, beside their function in development.

## Author Contributions

S.Z. performed experiments, analyzed data and wrote the paper. X.Z performed experiments. B. Q. and S.Z. conceived the study, designed the experiments. B.Q. supervised the study and wrote/edited the manuscript with S.Z.

## Acknowledgments

We thank the Caenorhabditis Genetics Center (CGC) (funded by NIH P40OD010440) for strains; Dr. Hong Zhang (Institute of Biophysics, Chinese Academy of Sciences) for providing strains of HZ3340 (*flag::let-363*); Dr. Xuna Wu of The Mass Spectrometry Facility in Yunnan University for MS. This work was supported by the Ministry of Science and Technology of the People’s Republic of China (2019YFA0803100, 2019YFA0802100), the National Natural Science Foundation of China (32170794), Yunnan Applied Basic Research Projects (202301AT070213, 202302AP370005, 202201AT070196), the Yunnan University Startup Program.

## Supplemental Figure Legends

**Figure S1.**
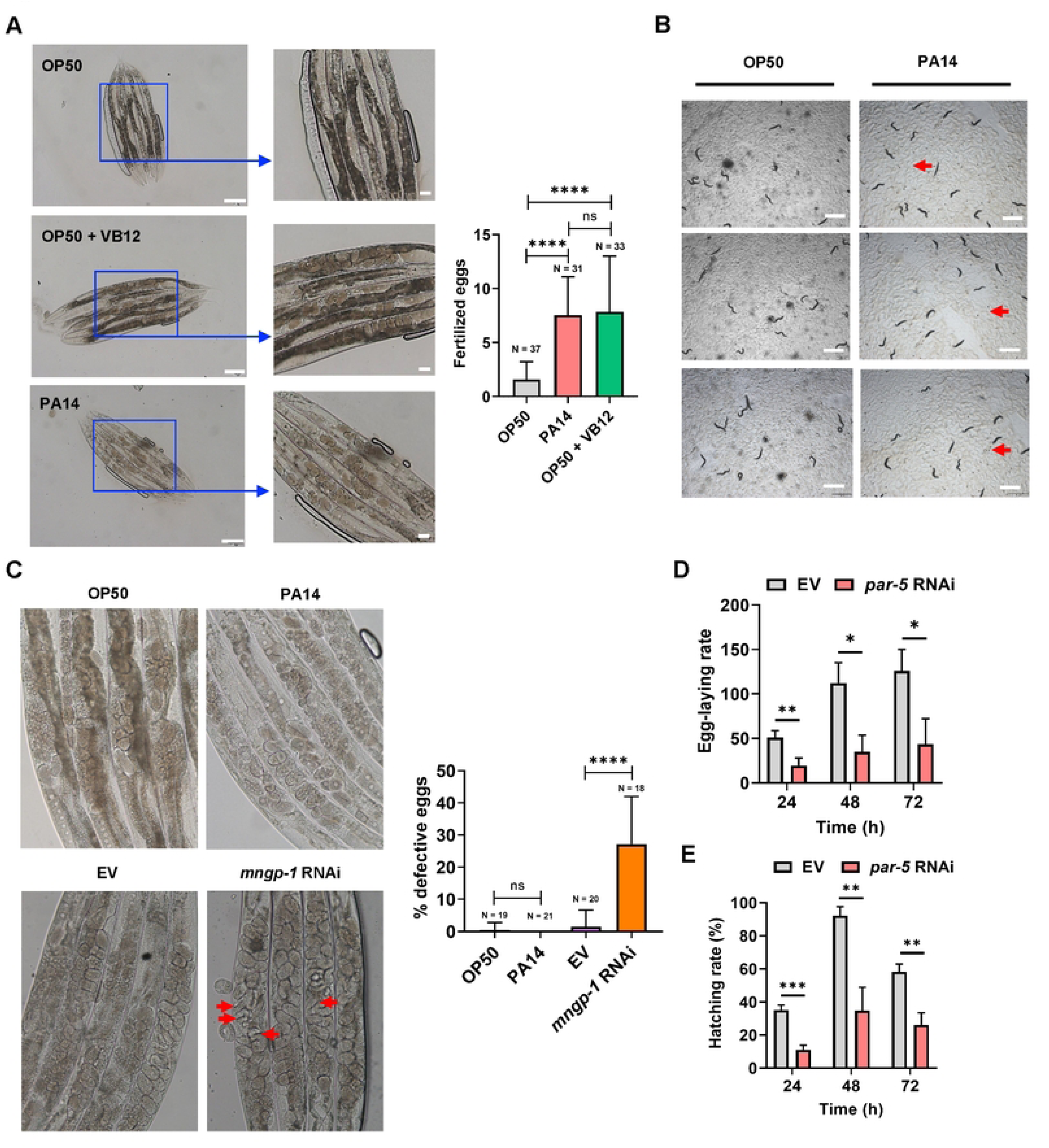
PAR-5 is critical for regulating egg-laying and hatching of worms. Related to Figure 1. (A) Microscope images and bar graph showing Vitamin B12 stimulates fertilized eggs formation. Scale bar, 100 μm.(B) Microscope images showing nematodes could lay and ovulate eggs normally under PA14 conditions, without forming any ‘worm bagging’, a phenomenon in which the eggs of nematodes hatch inside the parents’ body. Scale bar, 1000 μm (C) PA14 do not trigger the “bag of worms” phenotype. The red arrow indicates that *mngp-1* RNAi results in egg retention (the Bag phenotype) as positive control, consistent with previous studies (Fernández et al., 2022). Scale bar, 20 μm. (D) Bar graph showing RNAi inhibition of *par-5* reduces egg-laying rate of *C. elegans*. For each condition in assays, at least 5 animals were used. (E) Bar graph showing RNAi inhibition of *par-5* reduces hatching rate of *C. elegans.* For each condition in assays, at least 5 animals were used For all panels, N is the number of worms scored. Data are represented as mean ± SD. ∗∗∗∗p < 0.0001, ∗∗∗p < 0.001, ∗∗p < 0.01, ∗p < 0.05, ns, no significant difference. All data are representative of at least three independent experiments.

**Figure S2.**
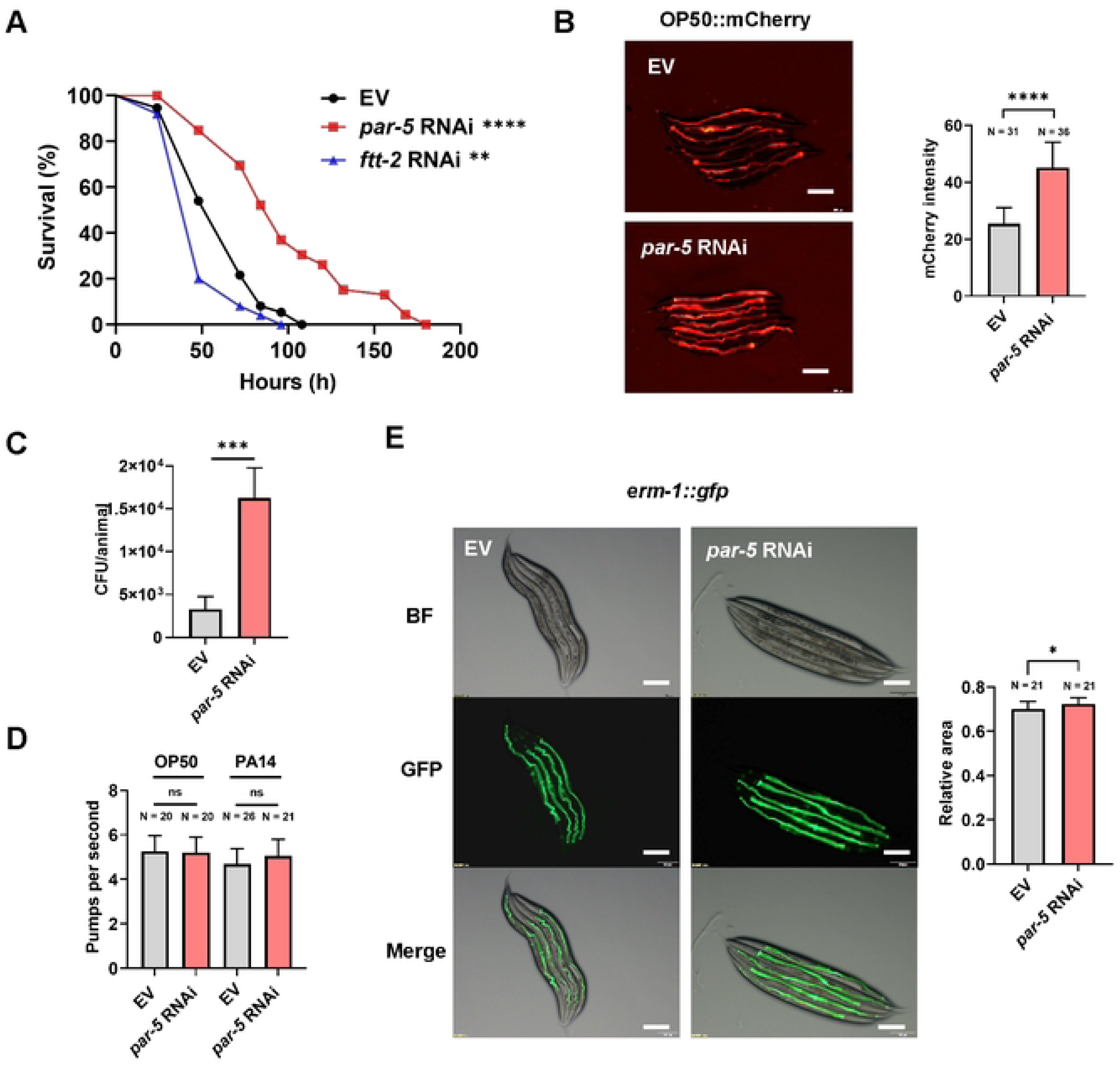
Summary of effects of PAR-5 on avoidance and clearance of pathogen. Related to Figure 2. (A) Knockdown of *ftt-2* by RNAi significantly enhances sensitivity to PA14 infection. *n = 50* worms for each sample. (B) Representative images and quantification of OP50-mCherry colonization in animals grown on control RNAi bacteria or *par-5* RNAi bacteria to L4 stage followed by exposure to OP50-mCherry for 24 h at 20°C. Scale bar, 200 μm. (C) Numbers of colony-forming units of OP50 were measured in worms subjected to *par-5* RNAi. These results are mean ± SD of four independent experiments performed in triplicate. (D) Bar graph showing pharyngeal pumping in animals grown on *E. coli* OP50 or PA14 fed with control RNAi bacteria or *par-5* RNAi bacteria. (E) Representative images and bar graph showing ERM-1::GFP localization in intestine, which was used to monitor the lumen morphogenesis. Scale bar, 200 μm. For all panels, N is the number of worms scored. Data are represented as mean ± SD. ∗∗∗∗p < 0.0001, ∗∗∗p < 0.001, ∗∗p < 0.01, ∗p < 0.05, ns, no significant difference. All data are representative of at least three independent experiments. The exact *P* values of statistics for all survival assays are listed in Table S1.

**Figure S3.**
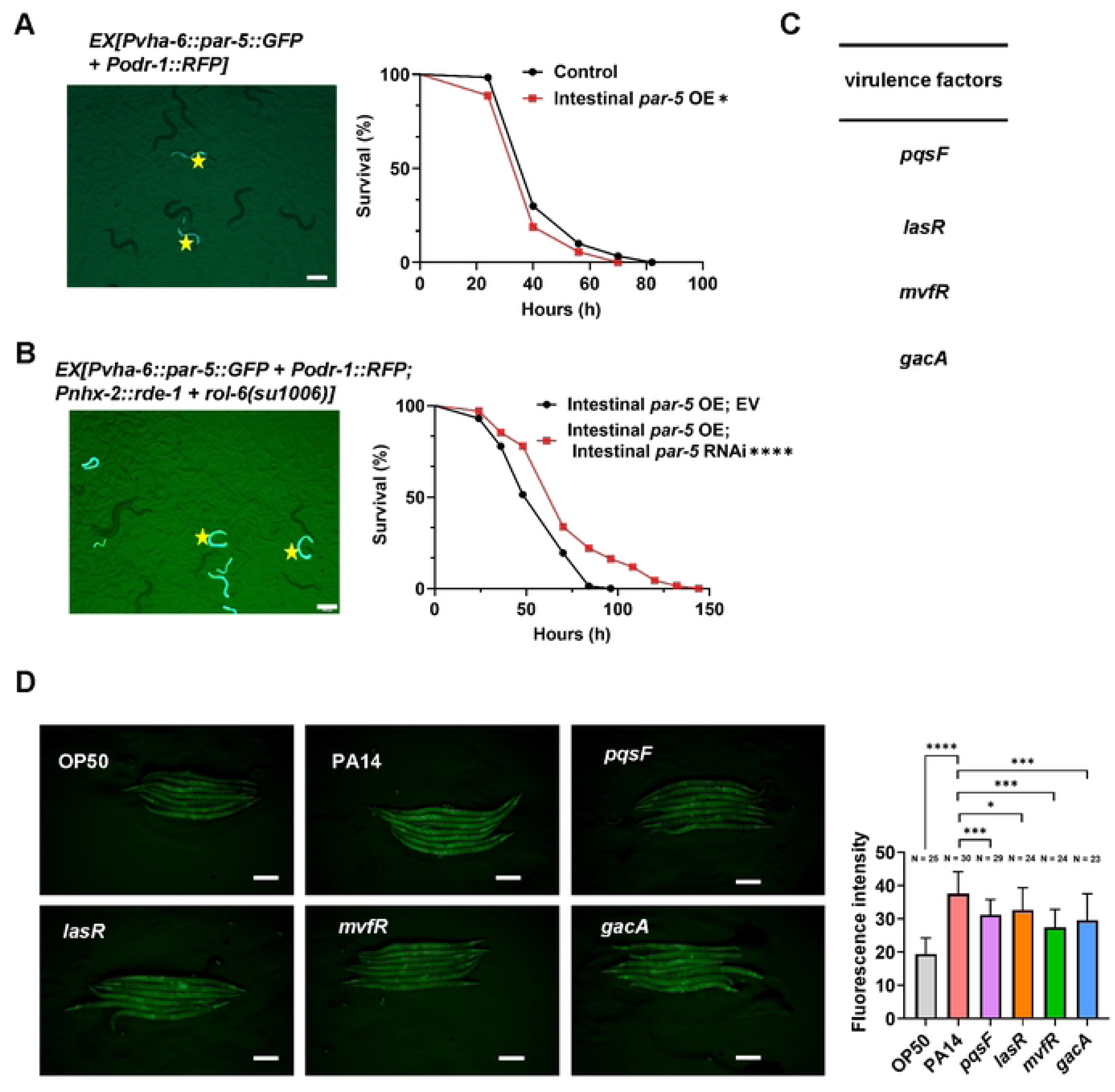
PAR-5 functions in the intestine, and is activated by virulence factors of PA14. Related to Figure 3. (A) Intestine-specific overexpression of PAR-5 decreases the resistance of worms to the PA14 infection. *n = 60* worms for each sample. Arrows indicate transgenic strains. Scale bar, 200 μm. (B) Intestinal *par-5* RNAi significantly enhances resistance to PA14 infection in animals with intestinal overexpression of *par-5*. *n = 75* worms for each sample. Arrows indicate transgenic strains. Scale bar, 200 μm. (C) List of PA14 mutants (virulence related genes) screened that may affect expression of *par-5*. (B) Microscope images and bar graph showing expression of *par-5::gfp* significantly decreases in L4 staged worms exposed to PA14 mutants (*pqsF*, *lasR*, *mvfR* and *gacA*) for 24 h at 20°C. Scale bar, 200 μm. For all panels, N is the number of worms scored. Data are represented as mean ± SD. ∗∗∗∗p < 0.0001, ∗∗∗p < 0.001, ∗∗p < 0.01, ∗p < 0.05, ns, no significant difference. All data are representative of at least three independent experiments.

**Figure S4.**
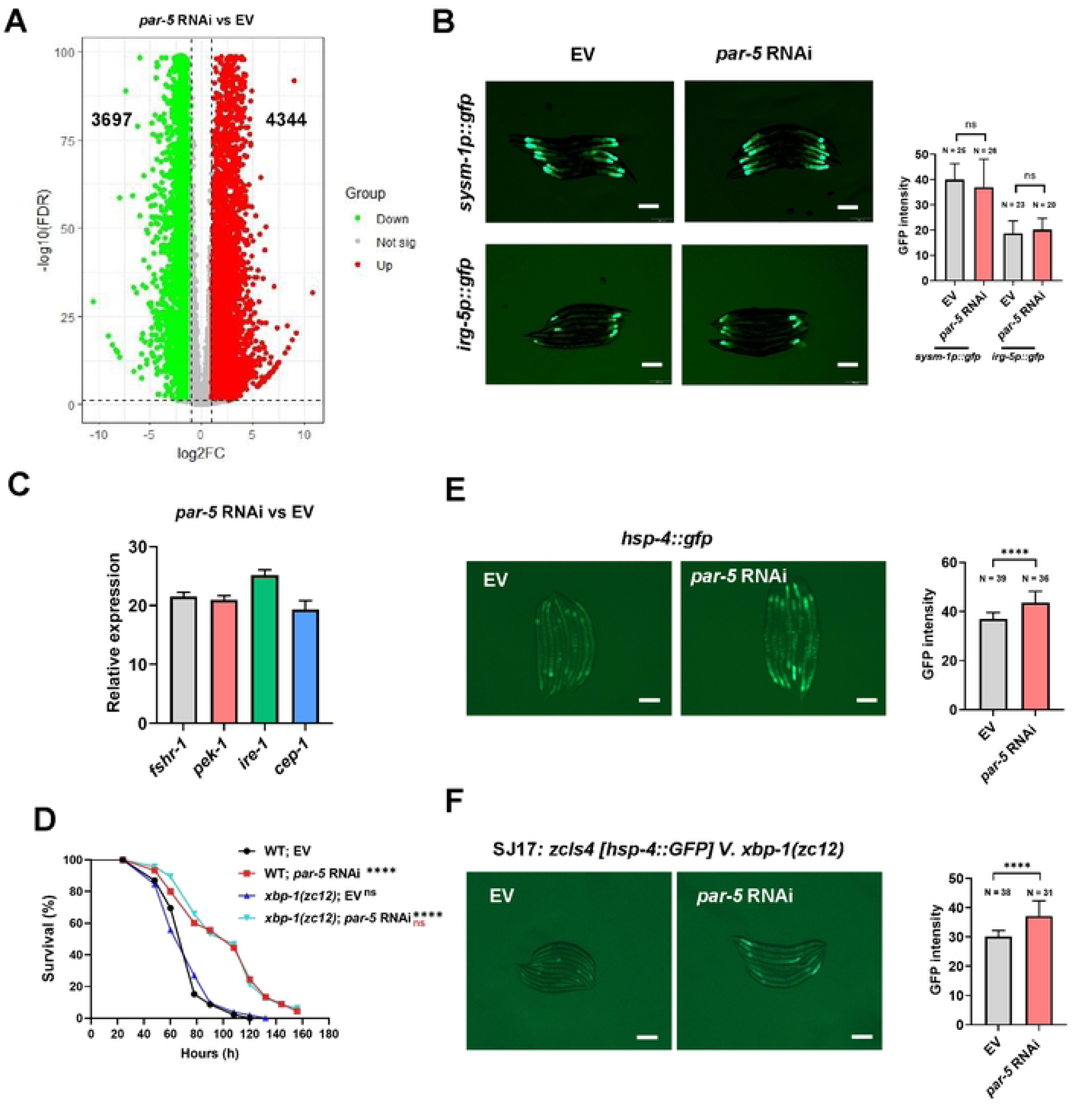
Summary of effects of PAR-5 on gene expression, immunity and UPR^ER^ of worms. Related to Figure 4. (A) Volcano plot of transcripts corresponding to differentially expressed genes in animals grown on PA14 with *par-5* RNAi versus vector control RNAi. Green and red colored dots indicate the differentially expressed genes that are increased or decreased, respectively. (B) Expression of *sysm-1p::gfp* and *irg-5p::gfp* in animals with *par-5* RNAi . Scale bar, 200 μm. (C) Comparison of differential expression of *fshr-1, pek-1, ire-1* and *cep-1* in *par-5* RNAi knock-down worms and vector control RNAi worms upon PA14 infection. (D) *xbp-1* mutation does not affect survival of worms with *par-5* RNAi upon PA14 infection. *n = 55* worms for each sample. (E) Microscope images and bar graph showing expression of *hsp-4::gfp* after *par-5* RNAi in N2 animals. Scale bar, 200 μm. (F) Microscope images and bar graph showing expression of *hsp-4::gfp* in *xbp-1(zc12*) mutants with *par-5* RNAi. Scale bar, 200 μm For all panels, N is the number of worms scored. Data are represented as mean ± SD. ∗∗∗∗p < 0.0001, ∗∗∗p < 0.001, ∗∗p < 0.01, ∗p < 0.05, ns, no significant difference. All data are representative of at least three independent experiments. The exact *P* values of statistics for all survival assays are listed in Table S1.

**Figure S5.**
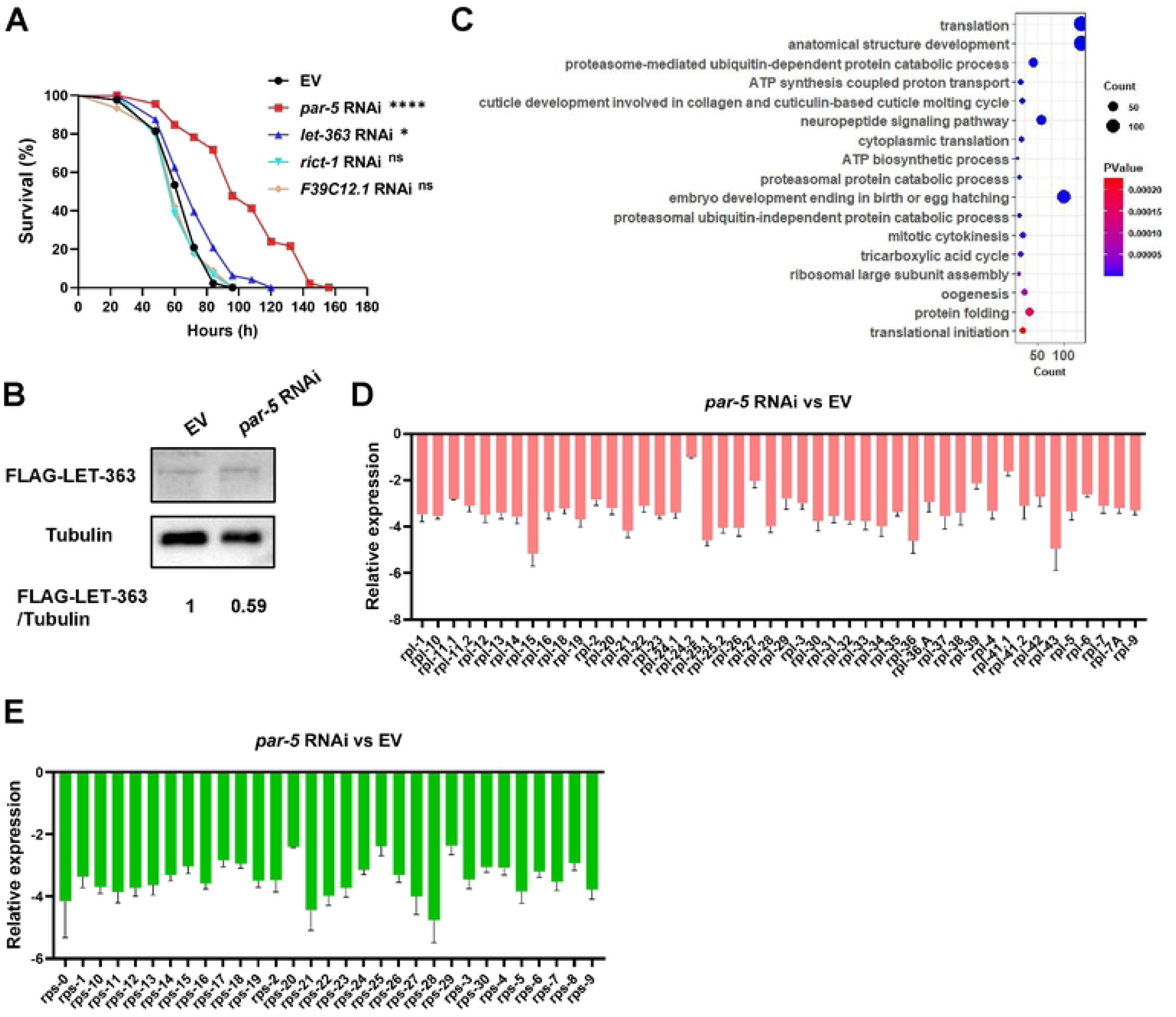
Summary of effects of PAR-5 on TOR activity and translation of worms. Related to Figure 5. (A) Knockdown of TOR signaling-related gene *let-363* by RNAi, but not F39C12.1 and *rict-1*, enhances resistance of N2 animals to PA14 infection. *n = 50* worms for each sample. (B) Western blot showing the level of LET-363 in animals fed with control RNAi bacteria or *par-5* RNAi bacteria. The blot is typical of three independent experiments. (C) GO enrichment of down-regulated gene in animals with *par-5* RNAi. The down-regulated genes are shown in Figure S4A and Table S5 (C and D) Comparison of differential expression of ribosomal protein subunit-related *rpl* genes (C) and *rps* genes (D) in *par-5* RNAi knock-down worms and vector control RNAi worms upon PA14 infection Data are represented as mean ± SD. ∗∗∗∗p < 0.0001, ∗∗∗p < 0.001, ∗∗p < 0.01, ∗p < 0.05, ns, no significant difference. All data are representative of at least three independent experiments. The exact *P* values of statistics for all survival assays are listed in Table S1.

**Figure S6.**
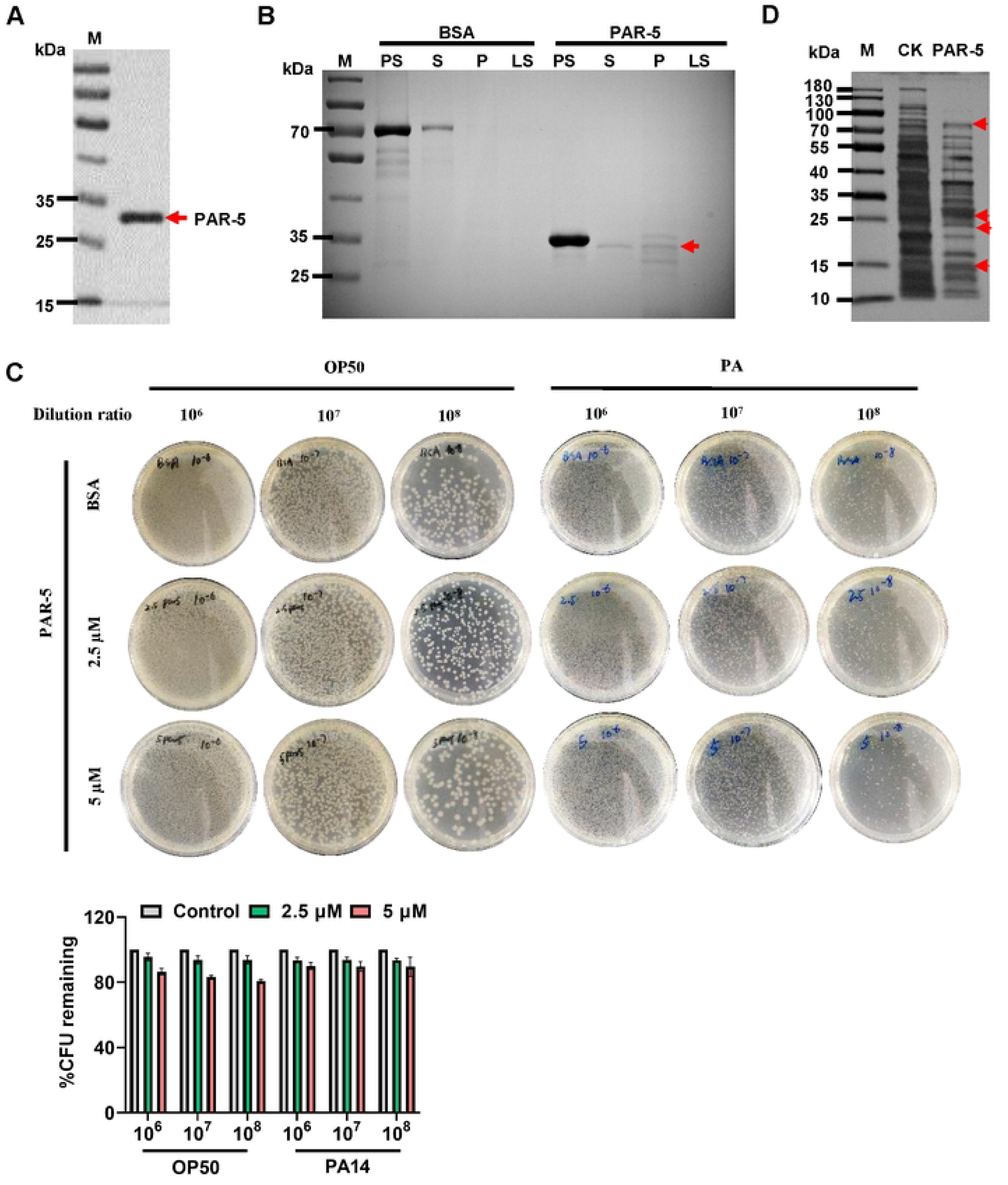
Summary of biochemical experiments of PAR-5 in vitro. Related to Figure 6. (A) Recombinant PAR-5 protein was expressed in *E. coli* BL21 and purified by nickel NTA-affinity chromatography. (B) *In vitro* bacterial binding assay shows that PGN binds PAR-5 protein purified from *E. coli* BL21 transfected with *his::par-5*. PGN was incubated with BSA (negative control) or His-PAR-5 protein, the PGN pellet was washed 3 times after incubation. Proteins associated with the washed pellet (P) and the incubated supernatant (S, LS) were detected by sliver-stained gel. M: marker; PS: purified protein (BSA or PAR-5); P: pellet; S: incubation supernatant; LS: supernatant of the last wash. (C) PAR-5 was added to ∼10^4^ CFU of log-phase *E. coli* OP50 or PA14 for 8 hours, and surviving bacteria were enumerated by dilution plating. (D) Sliver-stained gel showing the immunoprecipitation with purified His-PAR-5 against total isolated protein from PA14 Data are represented as mean ± SD. ∗∗∗∗p < 0.0001, ∗∗∗p < 0.001, ∗∗p < 0.01, ∗p < 0.05, ns, no significant difference. All data are representative of at least three independent experiments.

## Supplemental Tables

**Table S1.** Statistics for survival assay.

**Table S2.** Genes overlap among embryogenesis-related genes from studies published by Kamath et al (Kamath et al., 2003) and Sönnichsen et al (Sönnichsen et al., 2005), genes up-regulated by PA14 infection and RNA-seq data sets from the study published by Fletcher et al (Fletcher et al., 2019).

**Table S3.** Summary of up-regulated and down-regulated in animals with *par-5* RNAi under PA14 infection.

Table S4. GO enrichment analysis of the up-regulated genes identified in animals with *par-5* RNAi.

**Table S5.** GO enrichment analysis of the down-regulated genes identified in animals with *par-5* RNAi.

**Table S6.** Summary of upregulated and downregulated genes in PA14 with vs without PAR-5-treatment.

**Table S7.** Summary of down-regulated genes and PA14 virulence-related genes.

## Supplemental Data

**Data S1.** All unprocessed western blots.

**Data S2.** Survival Data for Replicate 2 and 3.

## Source Data. All source data is available in the supplemental information.

Source Data Fig. 1

Source Data Fig. 2

Source Data Fig. 3

Source Data Fig. 4

Source Data Fig. 5

Source Data Fig. 6

Source Data Fig. S1

Source Data Fig. S2

Source Data Fig. S3

Source Data Fig. S4

Source Data Fig. S5

Source Data Fig. S6

## KEY RESOURCES TABLE

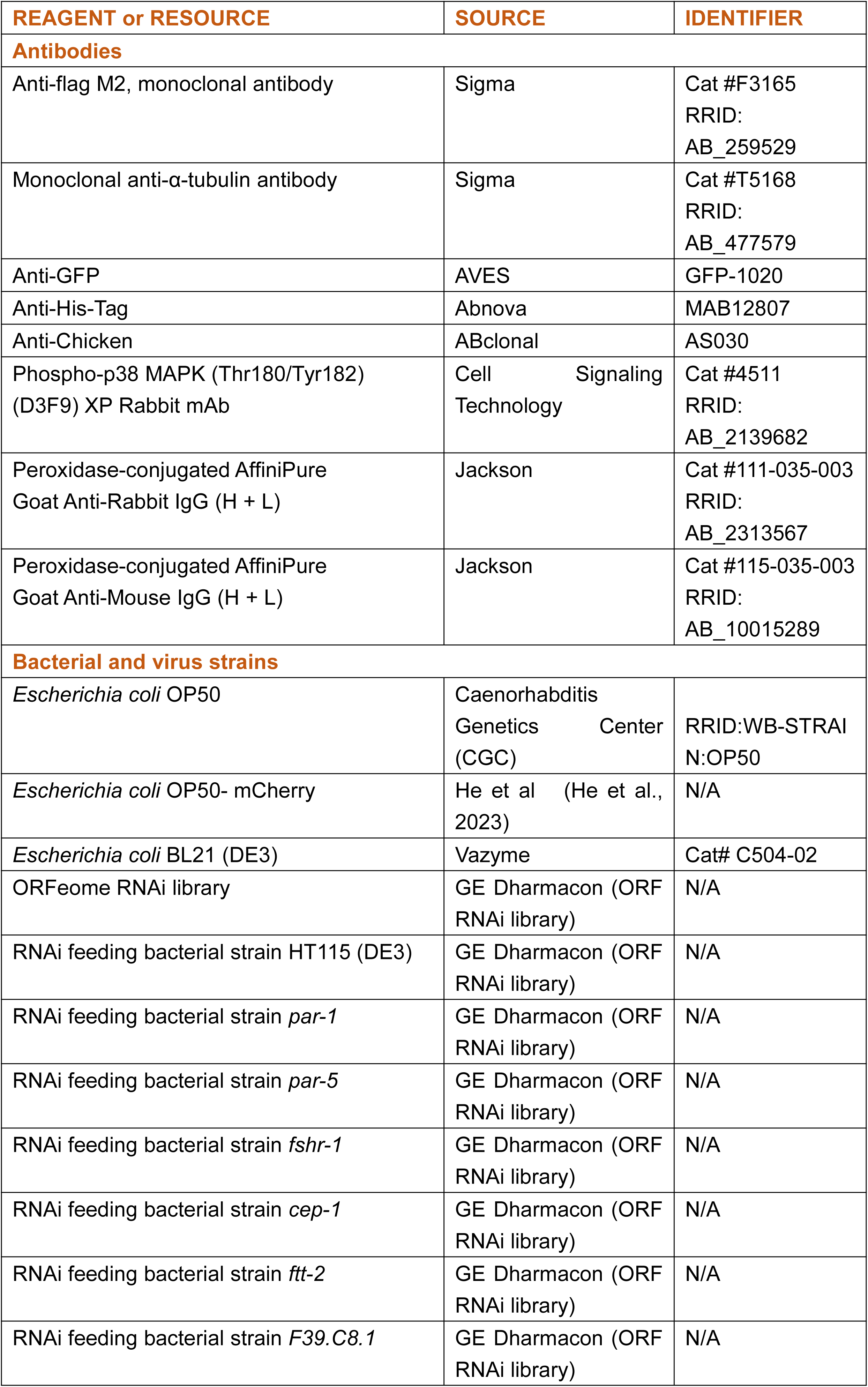
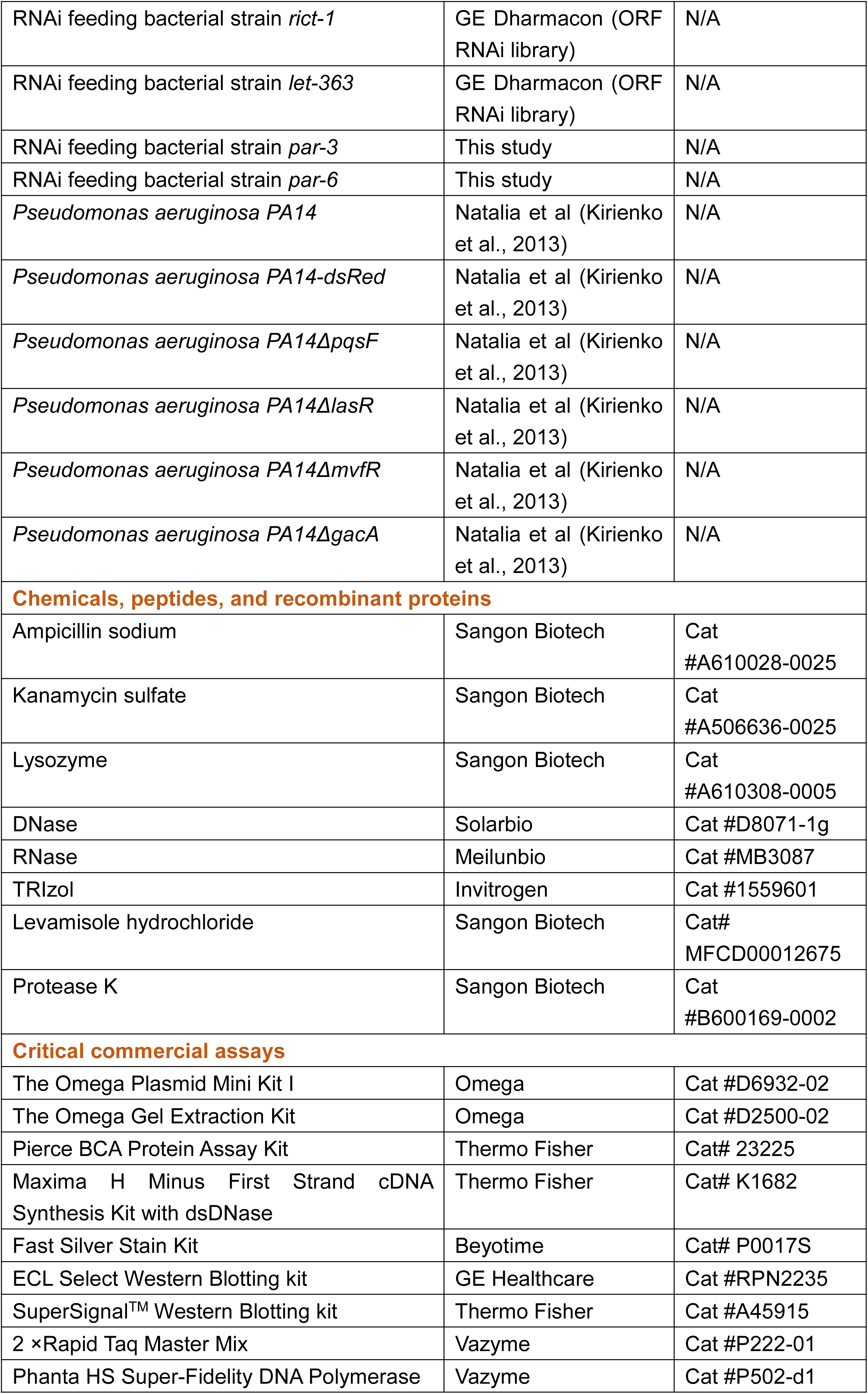
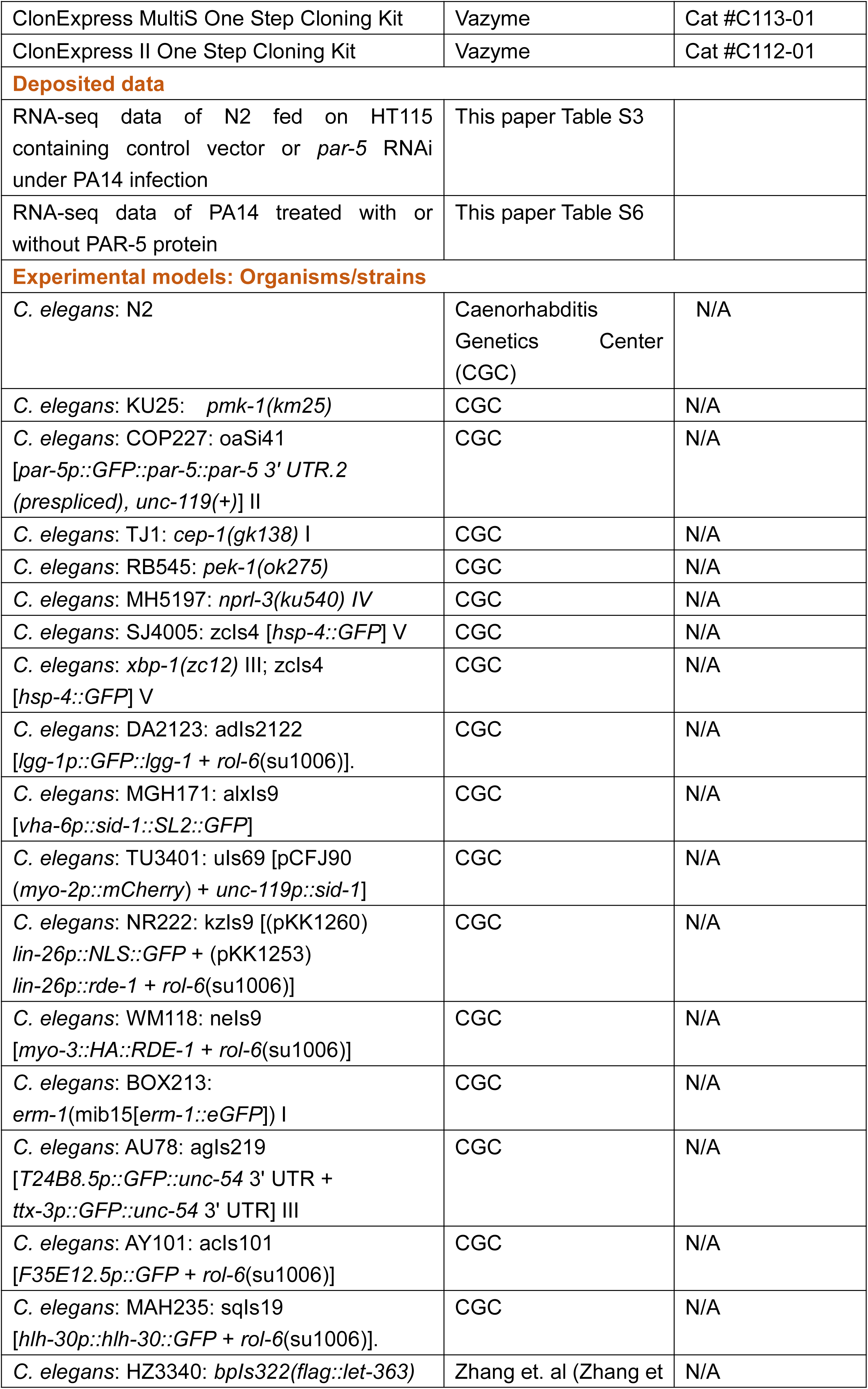
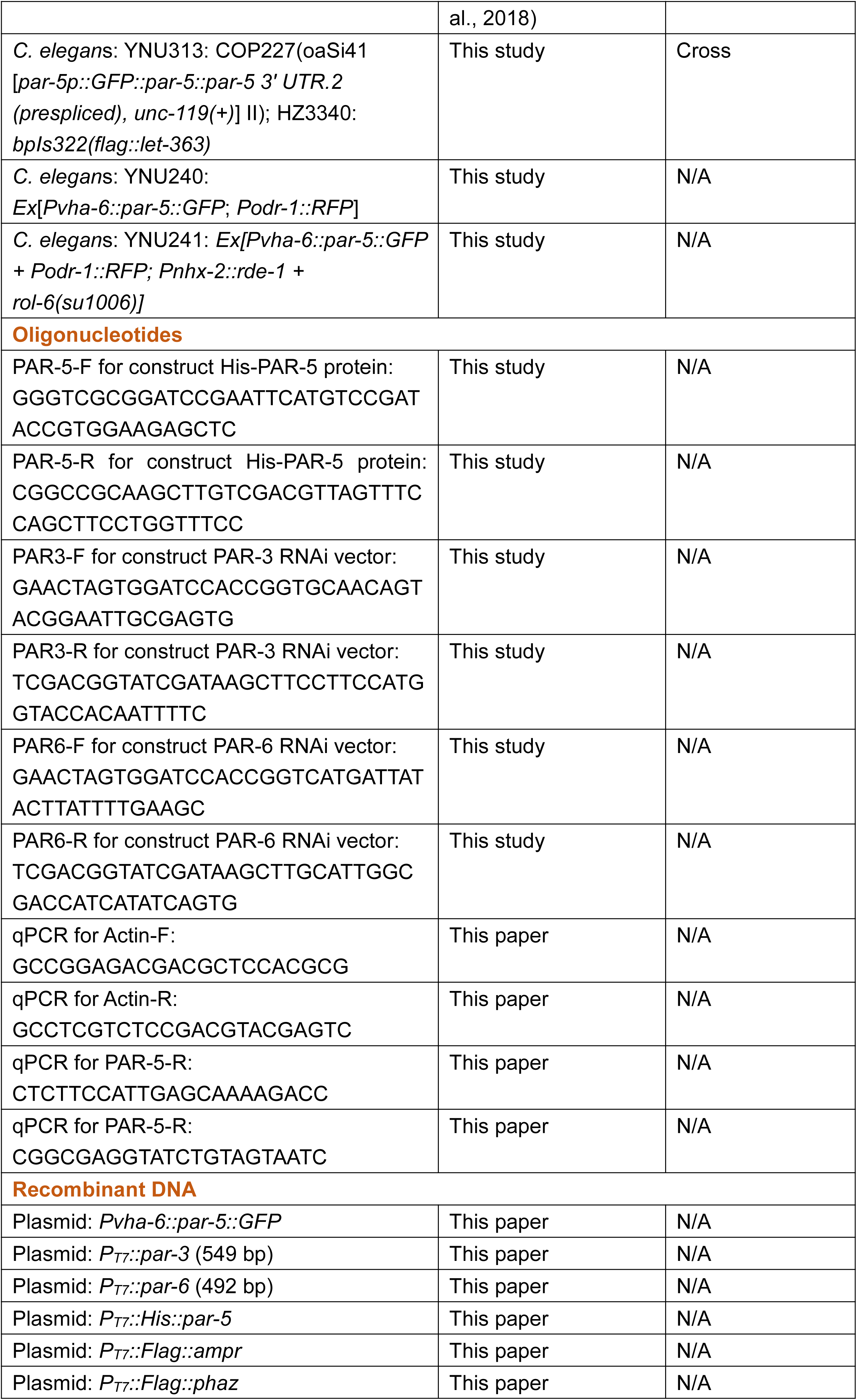
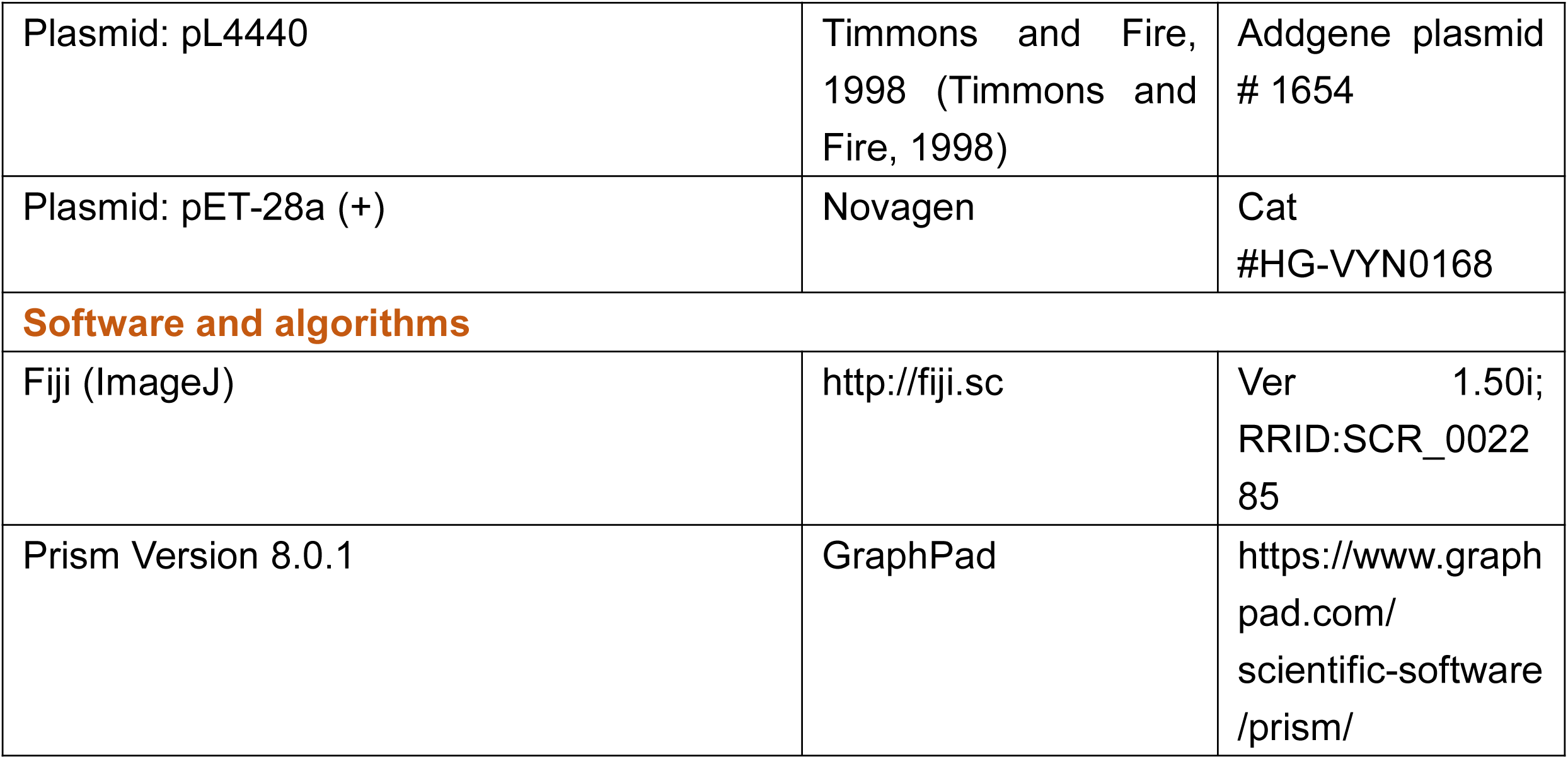

## STAR*METHODS

### RESOURCE AVAILABILITY

#### Lead contact

Further information and requests for reagents may be directed to the lead contact Bin Qi.

#### Materials availability

All reagents and strains generated by this study are available through request to the lead contact with a completed Material Transfer Agreement.

#### Data and code availability

RNA-seq data are accessible in Table S3 and Table S6. This paper does not report original code.

Any additional information required to reanalyze the data reported in this paper is available from the lead contact upon request.

### EXPERIMENTAL MODEL AND SUBJECT DETAILS

#### *C. elegans* strains and maintenance

Nematode stocks were maintained on nematode growth medium (NGM) plates seeded with bacteria (*E. coli* OP50) at 20℃.

1) The following strains/alleles were obtained from the Caenorhabditis Genetics Center (CGC) or as indicated: N2 Bristol (wildtype control strain); KU25: *pmk-1(km25)* IV; COP227: *oaSi41* [*par-5*p::*GFP*::*par-5*+ *unc-119*(+)] II; TJ1: *cep-1(gk138)* I; RB545: *pek-1(ok275)*; MH5197: *nprl-3(ku540)* IV;SJ4005: zcIs4 [*hsp-4::GFP*] V;; SJ17: *xbp-1(zc12)* III+ zcIs4 [*hsp-4::GFP*] V; MGH171: alxIs9 [*vha-6p::sid-1::SL2::GFP*]; VP303: kbIs7 [*nhx-2p::rde-1* + *rol-6*(su1006)]; TU3401: uIs69 [pCFJ90 (*myo-2p::mCherry*) + *unc-119p::sid-1*]; NR222: kzIs9 [(pKK1260) *lin-26p::NLS::GFP* + (pKK1253) *lin-26p::rde-1* + *rol-6*(su1006)]; WM118: neIs9 [*myo-3::HA::RDE-1* + *rol-6*(su1006)]; DA2123: adIs2122 [*lgg-1p::GFP::lgg-1* + *rol-6*(su1006)]; BOX213: *erm-1*(mib15[*erm-1::eGFP*]) I; AU78: agIs219 [*T24B8.5p::GFP::unc-54* 3’ UTR + *ttx-3p::GFP::unc-54* 3’ UTR] III; AY101: acIs101 [*F35E12.5p::GFP* + *rol-6*(su1006)]; MAH235: sqIs19 [*hlh-30p::hlh-30::GFP* + *rol-6*(su1006)]..

2) The following strains were obtained from Hong Zhang’s lab: HZ3340: bpIs322 (*flag::let-363*) (Zhang et al., 2018)

3) The following strains were constructed by this study:

YNU313: [*par-5p::GFP::par-5* + *unc-119*(+); *flag::let-363*] strain was constructed by crossing: COP227[*oaSi41* [*par-5*p::*GFP*::*par-5*+ *unc-119*(+)] with HZ3340[bpIs322 (*flag::let-363*)].

YNU240: [P*vha-6::par-5::GFP*; P*odr-1::RFP*] transgene strain was constructed by injecting plasmid P*vha-6::par-5::GFP* with P*odr-1::RFP* in N2 background.

YNU241: [P*vha-6::par-5::GFP* + P*odr-1::RFP*; P*nhx-2::rde-1* + *rol-6*(su1006)] transgene strain was constructed by injecting plasmid P*vha-6::par-5::GFP* with P*odr-1::RFP* in VP303 background.

#### Bacterial strains

1. *E. coli*-OP50, *E. coli*-OP50-mCherry, *Pseudomonas aeruginosa* PA14, *Pseudomonas aeruginosa* PA14 mutants, PA14-dsRed, *E. coli* HT115 were cultured at 37 ℃ in LB medium. A standard overnight cultured bacteria was then spread onto each Nematode growth media (NGM) plate.

### METHOD DETAILS

#### Generation of RNAi strains

To construct the RNAi plasmid, the gene segments was inserted into the pL4440 vector. PAR-3 (549 bp) was PCR-amplified from worm genome using oligonucleotides PAR3-F (5’-GAACTAGTGGATCCACCGGTGCAACAGTACGGAATTGCGAGTG-3’) and PAR3-R (5’-TCGACGGTATCGATAAGCTTCCTTCCATGGTACCACAATTTTC-3’), and was cloned into the corresponding sites (*Age* I/*Hin*d III) of pL4440 by ClonExpress II One Step Cloning Kit, which designated as *par-3* RNAi vector. PAR-6 (492 bp) was PCR-amplified from worm genome using oligonucleotides PAR6-F (5’-GAACTAGTGGATCCACCGGTCATGATTATACTTATTTTGAAGC-3’) and PAR6-R (5’-TCGACGGTATCGATAAGCTTGCATTGGCGACCATCATATCAGTG-3’), and was cloned into the corresponding sites (*Age* I/*Hin*d III) of pL4440 by ClonExpress II One Step Cloning Kit, which designated as *par-6* RNAi vector. The constructed plasmid was then transferred into *E. coli* HT115 (DE3) cells for expression.

#### Preparation of *E. coli*-OP50 bacteria

An overnight culture of *E. coli*-OP50 (37 ℃ in LB broth) was diluted into fresh LB broth (1:100 ratio). About 500 μl of the *E. coli*-OP50 was then spread onto each 6 cm Nematode growth media (NGM) plate when the diluted bacteria grew to OD_600_=0.5.

#### Generation of transgenes

1. To construct the *C. elegans* plasmid for expression of *par-5* in intestine, 1593 bp promoter of *vha-6* and genomic DNA of *par-5* was inserted into the pPD49.26 vector. DNA plasmid mixture containing P*vha-6::par-5::GFP* (10 ng/ul) and P*odr-1p:RFP* (25 ng/ul) was injected into the gonads of adult wild-type N2 animals.
2. The *C. elegans* plasmid for expression of *par-5* in intestine (P*vha-6::par-5::GFP*, 20 ng/ul) and P*odr-1p:RFP* (25 ng/ul) was injected into the gonads of adult VP303 animals that RNAi effective only in intestine.

#### Microscopy

For obversion/quantification of protein (GFP and mCherry) localization, larvae or adult worms were mounted on 3% agar pads in M9 buffer with 2.5 mM levamisole, and photographs were taken by using an inverted Leica STELLARIS 8 FALCON or Olympus BX53 microscope with a DP80 camera.

#### RNA interference (RNAi)

RNAi was used to generate loss-of-function RNAi phenotypes by feeding nematodes *E. coli* strain HT115(DE3) expressing double-stranded RNA (dsRNA) homologous to a target gene (Fraser et al., 2000). The *par-3* and *par-6* RNAi clones were constructed by this study. *E. coli* with the appropriate vectors were grown in LB broth containing ampicillin (100 μg/mL) at 37°C overnight and plated onto NGM plates containing 100 μg/mL ampicillin and 1 mM isopropyl β-D-thiogalactoside (IPTG) (RNAi plates).

L1 stage larvae were fed on RNAi expressing bacterial lawns and allowed to develop to L4 stage. L4 animals were then transferred to the NGM plate with PA14 bacteria for survival assay. Almost feeding RNAi experiments used bacterial clones from the MRC RNAi library (Kamath et al., 2003) or the ORF-RNAi Library (Rual et al., 2004).

#### *P. aeruginosa* lawn avoidance assays

The bacterial cultures were grown by inoculating individual bacterial colonies into 2 ml of LB broth and growing them on a shaker at 37°C overnight. Then, 20 µl of the culture was plated onto the center of 3.5 cm standard NGM plates. The plates were then incubated under the following conditions: 37°C for 12 hrs followed by 25°C for 12 hrs. Thirty synchronized L4 stage hermaphroditic animals were transferred inside the indicated bacterial lawns, and the numbers of animals off the lawns were counted at the specified times for each experiment. Three behaviors plates were used per trial in every experiment. The experiments were performed at 25°C. The percent occupancy was calculated as (N_on lawn_/N_total_) ×100. At least three independent experiments were performed.

#### Quantification of intestinal bacterial colonization

Thirty synchronized L4 stage animals grown on *E. coli* HT115 containing control vector or an RNAi clone targeting a gene were transferred into the indicated bacterial lawns (PA14-dsRed or *E. coli*-OP50-mCherry). L4 staged worms were fed at 25°C for the indicated times. Photographs were taken by using an inverted Leica STELLARIS 8 FALCON or Olympus BX53 microscope with a DP80 camera. The intensity of Fluorescence was calculated by Image J.

For measurement of PA14 or OP50 colony forming units (CFU), L4 staged worms were exposed to NGM plates with PA14-dsRed or *E. coli*-OP50-mCherry for 24 hours at 25°C. Then the animals were transferred from PA14-dsRed or *E. coli*-OP50-mCherry plates to the center of fresh empty plates for 10 min to eliminate PA14-dsRed or *E. coli*-OP50-mCherry stuck to their body. The worms were collected and soaked in M9 buffer containing 25 mM levamisole hydrochloride and 100 μg/ml ampicillin for 30 minutes at room temperature and washed three times with M9 buffer. Afterward, thirty worms per group were transferred into 50 μl PBS containing plus 0.01% Triton X-100 and ground. The lysates were diluted by 10-fold serial dilutions in sterilized water and seeded onto LB plates containing 100 μg/ml ampicillin to select for PA14-dsRed or *E. coli*-OP50-mCherry cells and grown overnight at 37°C. Single colonies were counted the next day and represented as the number of bacterial cells or CFU per animal. Three independent experiments were performed for each condition.

#### C. elegans killing assays on Pseudomonas aeruginosa PA14

Killing Assay was performed as previously described (Tan et al., 1999). Briefly, L4 stage larvae were manually transferred to 3.5 cm SK plates, for each strain, 3 plates of 30 worms were assayed. Worms were grown at 25°C and the surviving worms were counted every day. Alive worms were transferred to a fresh SK dish every 2-3 day during the reproductive period to eliminate self-progeny. Animals were scored at the times indicated and were considered dead when they failed to respond to touch. To prevent bacterial growth and rimmed with 150 μl of 10 mg/ml palmitic acid to prevent worms from crawling off the plates. At least three independent experiments were performed.

#### PAR-5::GFP induction assays

To measure the induction of PAR-5::GFP in *par-5p*::*par-5*::*GFP* reporter L4 stage animals, thirty synchronized L4 stage hermaphroditic animals grown on *E. coli* OP50 were transferred to the *P. aeruginosa* PA14 or PA14 mutants lawns for 24 hrs. These plates were incubated at 25°C for the indicated times, and then animals were prepared for fluorescence imaging. The fluorescence intensity in animal was quantified using ImageJ software.

#### qRT-PCR

Animals were synchronized by egg laying. Approximately 500 L1 stage animals (N2) were transferred to 6.5 cm RNAi plates seeded with *E. coli* HT115 containing control vector or RNAi clone targeting *par-5*. The animals were grown on the RNAi plates at 25°C until L4 stage. Then, the animals were transferred to 6.5 cm SK plates at 25°C for 12 hrs. The animals were collected, washed with M9 buffer. Total RNA was extracted using TRIzol reagent (Invitrogen) following manufacturer’s instructions. A total of 2 μg of RNA was reverse-transcribed with random primers using the Maxima H Minus First Strand cDNA Synthesis Kit with dsDNase (ThermoFisher, K1682). qPCR was performed by using PowerUp^TM^ SYBR^TM^ Green (ThermoFisher, A25742) on a real-time PCR machine (ABI QuantStudio I). Relative gene expression levels were calculated using the 2-ΔΔCt method. All samples were run in triplicate.

#### Pharyngeal pumping rates

L4 staged animal grown on HT115 containing control vector or *par-5* RNAi plates were used to measure the pharyngeal pumping rates. Pharyngeal pumping was counted during a 5 s interval using Olympus MVX10 dissecting microscope with a DP80 camera. Then calculated to per second.

#### Worm total protein extraction

Worms were lysed by freeze-thawing five times in protein lysis buffer containing PMSF in liquid nitrogen. The worms were then ground in a tissue grinding tube. The resultant liquid was then centrifuged at 15000 rpm for 30 min at 4℃, and the supernatant was taken as the total protein of worms. The protein concentration was measured by using the Pierce BCA protein assay kit (ThermoFisher, 23227).

#### Protein purification

Protein purification was done with three biological replicates that were protein expression, collected, and purification.

(1) Protein expression

PAR-5 gene was cloned into a PT7 vector (P_T7_-His-PAR-5) and heterologously expressed in *E. coli* OP50. The cDNA encoding complete *par-5* was amplified by PCR using oligonucleotides PAR-F

(5’-GGGTCGCGGATCCGAATTCATGTCCGATACCGTGGAAGAGCTC-3’) and PAR-R (5’-CGGCCGCAAGCTTGTCGACGTTAGTTTCCAGCTTCCTGGTTTCC-3’). PAR-5 was cloned into the corresponding sites (*Eco*R I/*Sal* I) of pET-28a (+) by ClonExpress II One Step Cloning Kit. Bacteria was grown in LB broth containing ampicillin (100 μg/ml) at 37°C overnight. Inoculating individual bacterial colonies into 10 mL of LB broth containing ampicillin (100 μg/mL), and growing them overnight on a shaker at 37°C. Expand the volume of LB broth for culture bacteria at 1:100, after 2-3 hrs (OD_600_=0.6-0.8) add 0.4 mM isopropyl β-D-thiogalactoside (IPTG) into culture medium to induce PAR-5 expression.

(2) Collecting and cleaving

Bacteria was pelleted by centrifugation at 8000 rpm for 20 min. Harvested bacteria was re-suspended in the buffer-A, and lysed by sonication at 4℃.

(3) Protein purification

PAR-5 protein was expressed in the *E. coli* strain BL21(DE3) with a N-terminal His-tag. The soluble fraction of bacteria was purified using a glutathione sepharose column (GE Healthcare, 17-0756-01). The proteins were then affinity purified using a Ni2^+^ Sepharose column (GE Healthcare, 17-5268-01) and eluted from the column with 200 mM imidazole. Purified proteins were concentrated using 10 kDa molecular mass cut-off centrifugal filter units to approximately 200 ng μl^−1^ final concentration and dialysed twice using a dialysis buffer containing 20 mM Tris-HCl (pH 8.0), 300 mM NaCl and 5% (v/v) glycerol at 4 °C for 6 hrs with magnetic stirring. Insoluble aggregates after dialysis were removed by high-speed centrifugation. The proteins were then diluted to 100 ng μl^−1^ final concentration with the dialysis buffer and stored at −80 °C in aliquots. The concentrations of purified proteins were determined by Pierce BCA Protein Assay Kit (Thermo Fisher, 23225).

#### Protein-bacterial binding assay

*P. aeruginosa* PA14 was cultured in LB at 37 ℃ to OD_600_ values. 2 ml of bacteria was centrifuged at 8000 rpm for 1 min, the supernatant was discarded, and the pellet was washed with PBS buffer three times and finally re-suspend in PBS buffer.

This 500 μl bacterial suspension was incubated with the same amount of protein and incubated at 4 ℃ with rotation overnight. At the end of incubation, the bacteria were pelleted, the incubated supernatant was removed, and the pellet was washed three times with PBS buffer. Proteins associated with the washed pellet and the incubated supernatant were detected by Western blot.

#### IP-MS

For *C. elegans*, 5 mg of total protein from COP227 (*par-5p::gfp::par-5*) worm lysate was mixed with Anti-GFP beads (Abmart, M20016M) (30 μl) at 4℃ overnight and then washed five times with PBS wash buffer. Proteins were eluted using 1% SDS by boiling and detected by silver staining and Western blot analysis probed with anti-GFP chicken mAb (AVES, AB_2307313), then subjected to mass spectrometry analysis.

For *P. aeruginosa* PA14, mixture of lysate of an overnight culture of *P. aeruginosa* PA14 and His-PAR-5 protein was mixed with Ni-NTA agarose (QIAGEN, 30210) (30 μl) at 4℃ overnight and then washed five times with PBS wash buffer. Proteins were eluted using 1% SDS by boiling and detected by silver staining and Western blot analysis probed with Anti-His-Tag mAb (Abnova, MAB12807), then subjected to mass spectrometry analysis.

#### Co-immunoprecipitation

For proteins in *C. elegans*, lysate of transgenic worms YNU-313 (carrying *par-5p::GFP::par-5* and *flag::let-363*) was mixed with Anti-GFP beads (Abmart, M20016M) or Anti-FLAG beads (Sigma, A2220) (30 μl) at 4℃ overnight and then washed five times with PBS wash buffer. Proteins were eluted using 1% SDS by boiling and detected by Western blot analysis probed with Anti-GFP chicken mAb (AVES, AB_2307313) and anti-FLAG mouse mAb (Sigma, F3165).

For proteins in *P. aeruginosa* PA14, the mixture of lysate of *E. coli* BL21 expressing recombinant Flag-tagged virulence-related protein and His-PAR-5 protein was incubated with anti-FLAG beads (Sigma, A2220) (30 μl) at 4℃ overnight and then washed five times with PBS wash buffer. Proteins were eluted using 1% SDS by boiling and detected by Western blot analysis probed with anti-His-Tag mAb (Abnova, MAB12807) and Anti-FLAG mouse mAb (Sigma, F3165).

#### Western blot

The targeting proteins were analyzed by standard Western blot methods and probed with following antibodies: anti-His (dilution = 1:10000, Abnova, MAB12807), anti-FLAG (dilution = 1:5000, Sigma, F3165), Monoclonal anti-a-tubulin antibody (dilution = 1:10000, Sigma, T5168), anti-GFP (dilution = 1:5000; AVES, GFP-1020), anti-Chicken IgY (dilution = 1:5000; ABclonal, AS030), anti-p-p38 (dilution = 1:5000; Cell Signaling, 4511S), Peroxidase-conjugated AffiniPure Goat Anti-Rabbit IgG (H + L) (dilution = 1:5000, Jackson, 111-035-003), Peroxidase-conjugated AffiniPure Goat Anti-Mouse IgG (H + L) (dilution = 1:10000, Jackson, 115-035-003).

#### Preparation of samples for RNA sequencing

RNA-seq was done with three biological replicates that were independently generated, collected, and processed.

For the pair-I (*E. coli* OP50 vs *P. aeruginosa* PA14) sample: L4 stage wild type (N2) worms growing on *E. coli* OP50 plates were transfer to the *E. coli* OP50 or *P. aeruginosa* PA14, the samples were collected after 10 hours at 25°C.

For the pair-II (*E. coli* HT115 vs *par-5* RNAi) sample: L4 stage wild type (N2) worms growing on *E. coli* HT115 and *par-5* RNAi plates were transfer to the *P. aeruginosa* PA14, the samples were collected after 10 hours at 25°C.

For the pair-III (*P. aeruginosa* PA14 vs PAR-5-treated *P. aeruginosa* PA14) sample: An overnight culture of *P. aeruginosa* PA14 (37℃ in LB broth) was diluted into fresh LB broth. Diluted bacterial solution were cultured in LB broth adding purified protein (BSA or PAR-5) for 14 hrs at 37℃.

#### RNA sequencing and data processing

For *C. elegans*, a total amount of 1 μg RNA per sample was used as input material for the RNA sample preparations. Sequencing libraries were generated using NEBNext UltraTM RNA Library Prep Kit for Illumina (NEB, USA) following manufacturer’s recommendations and index codes were added to attribute sequences to each sample. The library preparations were sequenced on an Illumina platform and paired-end reads were generated. The raw reads were further processed with a bioinformatic pipeline tool, BMKCloud (www.biocloud.net) online platform.

For *P. aeruginosa* PA14, a total amount of 1 μg RNA per sample was used as input material for the RNA sample preparations. RNA samples were sent to Genewiz for further processing, including rRNA depletion and dUTP incorporation for strand-specific sequencing, and sequenced with an Illumina HiSeq 2500 platform. Six fastq files were mapped to the reference PA14 genome using Bowtie2 (Langmead and Salzberg, 2012) with an ∼97% success rate to generate SAM (sequence alignment map) files. SAM files containing ∼2 × 108 reads in total were merged, sorted, and indexed with SAMtools (Li et al., 2009). Read coverage was visualized with Integrative Genomics Viewer (IGV) software (Robinson et al., 2011).

Differential expression analysis of two conditions/groups was performed using the DESeq2. Gene Ontology (GO) enrichment analysis of the differentially expressed genes (DEGs) was implemented by the GOseq R packages based Wallenius non-central hyper-geometric distribution, which can adjust for gene length bias in DEGs. RNA-seq data are available in Table S3 and Table S6.

#### Pyocyanin quantitation assay

The pyocyanin assay is based upon the absorbance of excreted pyocyanin at 520 nm in acidic solution (Essar et al., 1990). Overnight cultured *P. aeruginosa* PA14 was diluted into 5ml fresh medium (1:1000) mixed with 200 μM PAR-5 protein following 14 h of aerobic growth with shaking at 37 °C. Cells were separated from culture fluids via centrifugation at 15000 g for 15 min. A 3 ml supernatant from above culture of PAR-5-treated *P. aeruginosa* PA14 was mixed with 1.8 ml of chloroform. The pyocyanin from the chloroform phase was then extracted into 0.6 ml of 0.2 N HCl, giving it a pink to deep red color, indicating the presence of pyocyanin. The absorbance was measured at 520 nm. Concentrations, expressed as micrograms of pyocyanin produced per milliliter of culture supernatant, were determined by multiplying the optical density at 520 nm (OD_520_) by 17.072.

#### Analysis of autophagic events using an LGG-1 reporter strain

The level of autophagy in various mutants was assessed using an LGG-1::GFP translational reporter characterized previously (Melendez et al., 2003). GFP-positive puncta were counted (using Olympus BX53 microscope with a DP80 camera) in randomly selected seam cells of L3 stage larvae and calculating the average number of foci per cell. When performing RNAi experiments to count LGG-1::GFP-positive foci, L1 stage animals were fed the RNAi bacteria at 25℃, and the L3 progeny of their progeny (“F2 generation”) were examined. A number of seam cells randomly selected from at least 10 animals in each experiment for photo recording and analysis at least three independent trials and averaged.

#### HLH-30 nuclear translocation

The nuclear localization of HLH-30::GFP was visualized by using the Olympus BX53 microscope with a DP80 camera. About 20 animals were used per condition, and the percentage of animals showing the nuclear translocation was counted.

#### Analysis of Bag phenotype

To analyze the Bag phenotype, a number of L4 stage hermaphrodites (more than 100 worms) grown on *E. coli* OP50 or RNAi bacteria plates (RNAi control or *nmgp-1* RNAi) were transfer to the fresh plates containing the same culture medium or *P. aeruginosa* PA14. After laying eggs, the total number of fertilized eggs within the insect body and the number of late embryos inside the uterus (“bag-of-worms”) were observed and counted (using Olympus BX53 microscope with a DP80 camera) in approximately 20 randomly selected nematodes at indicated time (in Figure S1C). Defective eggs Rate = N_late embryos inside the uterus_ /N_fertilized eggs within the insect body_ × 100%.

#### Measurement of early formation of fertilized eggs

To analyze early zygotes production, a number of L4 stage hermaphrodites (more than 100 worms) grown on *E. coli* OP50 (with or without 1 μM Vitamin B12) or RNAi bacteria plates (RNAi control or *par-5* RNAi) were transfer to the *E. coli* OP50 or *P. aeruginosa* PA14 for 7-18 hrs at 25 ℃ . The number of fertilized eggs formed were observed and counted (using Olympus BX53 microscope with a DP80 camera) in at least 40 randomly selected nematodes at indicated time (in Figure 1A, 1D and S1A) .

#### Analysis of the effect of PAR-5 on the oviposition cycle

Five young adult stage animals grown on RNAi control or *par-*5 RNAi plates were transferred to a new *E. coli* OP50 culture plate, and the number of eggs were counted (using Olympus BX53 microscope with a DP80 camera) on the plate after the transfer of worms to the plate once a day until the worms does not lay eggs (N_laid egg_ = N_egg per dish_/N_animals_; Hatching Rate = (N_hatching egg_/N_animals_)/N_laid egg_×100%; N_zygote_=SUM_zygote per dish_/N_animals_×100%).

### QUANTIFICATION AND STATISTICAL ANALYSIS

#### Quantification

ImageJ software was used for quantifying relative fluorescence intensity of PAR-5::GFP reporter or PA14-dsRed.

#### Statistical analysis

All statistical analyses were preformed using Student’s t-test, except for survival assays which were analyzed by using Log-rank (Mantel-Cox) test and Figures S3D which were analyzed by using One-way ANOVA.

Two-tailed unpaired t test was used for statistical analysis of two groups of samples. Data are presented as Mean ± SD, and p < 0.05 was considered a significant difference.

